# Calcium Sensitive Allostery Regulates the PI(4,5)P2 Binding Site of the Dysferlin C2A Domain

**DOI:** 10.1101/2021.02.10.430549

**Authors:** Shauna C. Otto, Patrick N. Reardon, Tanushri M. Kumar, Chapman J. Kuykendall, Colin P. Johnson

## Abstract

C2 domains are the second-most abundant calcium binding module in the proteome. Activity of the muscular dystrophy associated protein dysferlin is dependent on the C2A domain at the N-terminus of the protein, which couples calcium and PI(4,5)P2 binding through an unknown mechanism. Using solution state nuclear magnetic resonance spectroscopy we confirm the phosphoinositide binding site for the domain and find that calcium binding attenuates millisecond to microsecond motions at both in the calcium binding loops and the concave face of the C2A, including a portion of the phosphoinositide binding site. Our results support a model whereby increasing calcium concentrations shift the phosphoinositide binding pocket of C2A into a binding-competent state, allowing for calcium dependent membrane targeting. This model contrasts with the canonical mechanism for C2 domain-phosphoinositide interaction and provides a basis for how pathogenic mutations in the C2A domain result in loss of function and disease.

## Introduction

C2 domains are lipid-binding modules that can integrate calcium and lipid signals (1, 2). Although sequence similarity can be quite low between C2 domains, the tertiary structure of the 8-stranded β-sandwich core of the domain is typically conserved and unaffected by calcium (3, 4). By contrast, the structure of the calcium binding loops at the end of C2 domains are diverse, tune the domain’s sensitivity to calcium, and facilitate binding to anionic or zwitterionic lipid headgroups (5, 6). Separate from this calcium-lipid binding pocket, some C2 domains also contain a conserved swath of basic and accessory residues along the concave side of the domain, the polybasic cluster, binds phosphatidylinositol 4,5- bisphosphate (PI(4,5)P2) and other phosphoinositide species (7–10). A causal link between calcium and phosphoinositide binding has been noted for several of these C2 domains (11– 14) but a universal structural one has yet to be defined.

The muscle protein dysferlin contains seven predicted C2 domains (Figure 1), but only the N-terminal C2A is reported to bind both PI(4,5)P2(13) and calcium (6, 13). Cell-based assays have confirmed the importance of this domain for dysferlin function (15, 16), with either mutation or domain truncation resulting in functional deficiency and loss of membrane localization. For example, the muscular dystrophy-associated missense mutation V67D within the C2A do-main results in loss of colocalization between dysferlin and PI(4,5)P2 (17). Dysferlin and PI(4,5)P2 interact in muscle in a number of cellular contexts (Figure 1). First, imaging techniques have unveiled a pool of dysferlin within the transverse tubule system (18), a structure highly enriched in PI(4,5)P2 (19). Dysferlin interacts closely with dihydropyridine receptors (20) at the triad junction and loss of dysferlin activity results in defects in Ca^2+^ homeostasis within muscle (21). Dysferlin has been shown to be required for the development and maintenance of the transverse tubule system, a process which also requires the presence of PI(4,5)P2. Strikingly when overexpressed dysferlin can induce the formation of extensive transverse-tubule-like structures in non-muscle cells in a PI(4,5)P2-dependant manner (17). Second, repair molecules, including PI(4,5)P2 and dysferlin, accumulate at muscle cell membrane tears, and that dysferlin confers calcium sensitivity to membrane fusion events that mediate repair (22). Failure to repair these mechanically-induced membrane tears are responsible for several human pathologies (23).

**Fig. 1.**
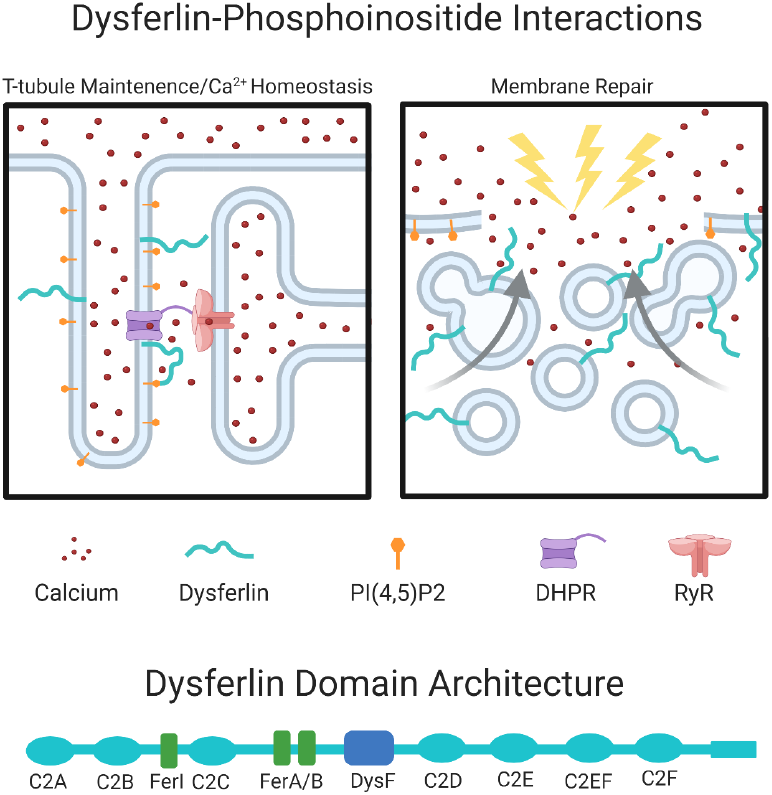
Dysferlin-phosphoinositide interactions. (Top) Dysferlin and PI(4,5)P2 are both found in the transverse tubule system, along with the dihydropyridine (DHPR) and ryanodine (RyR) receptors. PI(4,5)P2 and dysferlin cluster at wound sites in response to calcium. (Bottom) Dysferlin contains 7 C2 domains. Created with BioRender.com.

How the C2A domain coordinates calcium signals with PI(4,5)P2 binding is unknown, and a deeper understanding of the mechanisms that govern this process is essential to develop therapeutic strategies for dysferlinopathies. In this study we apply solution NMR spectroscopy to directly monitor the dynamics of the C2A domain under the influence of these effectors, and map the binding face of the domain that interacts with the headgroup of PI(4,5)P2, inositol 1,4,5-triphosphate (IP3). We find that binding of calcium restricts motion within the apical end of the domain, especially the calcium binding loops, as well as residues involved in PI(4,5)P2 binding. In addition, IP3 binding further restricts calcium binding loop movement, suggesting that both PI(4,5)P2 and calcium cooperate to influence domain structure and membrane association. The restriction of apical end dynamics suggests that calcium organizes the top half of the phosphoinositide binding region prior to membrane interaction in a way that may be representative of other calcium-sensitive phosphoinositide binding C2 domains.

## Results

### Resonance Assignments and Secondary Structure

To structurally probe dysferlin C2A, we generated a recombinant form of the domain composed of residues 1-129 with a hexhistidine tag at the N-terminus. The [^1^H-^15^N]-TROSY spectra was found to be well dispersed, indicating that the domain is folded (Figure 2A). We assigned 108 of the 120 non-proline residues, and of the unassigned residues, 75% (9 out of 12) were in the loop regions between β-strands. Of the remaining three residues two were located next to the C-terminus and one is adjacent to calcium binding loop 3 (CBL3) (Figure 2B). As shown in Figure S1, secondary structure prediction based on Talos+ showed good agreement with the existing crystal structure of the dysferlin C2A domain (PDB: 4IHB) (24) and the recently reported solution structure (PDB: 7KRB) (25). This indicates that the structure of the protein in solution is similar to that observed in the crystal structure, under calcium saturating conditions.

**Fig. 2.**
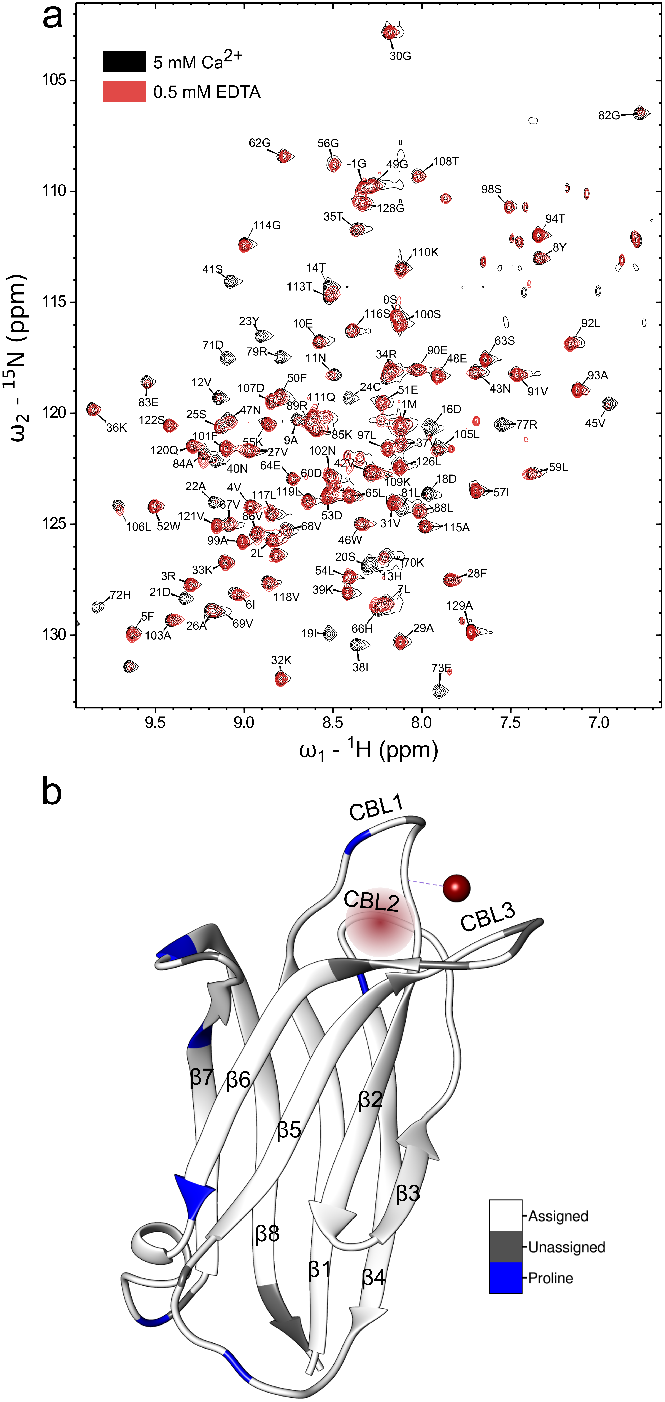
a) [^1^H-^15^N]-TROSY spectrum of human Dysferlin C2A (1-129) in the presence (black, 5 mM CaCl_2_) or absence (red, 0.5 mM EDTA) of calcium. b) assignment completion mapped onto the structure (4IHB). β-strands 1-8 and calcium binding loops (CBLs) 1-3 are labeled. Density for only a single calcium was seen in the crystal structure (solid red sphere), the predicted second calcium binding site is indicated by the diffuse red sphere.

### Calcium Reduces the Flexibility of the C2 Domain

To first gain insight into the influence of calcium on the structure of C2A, we collected [^1^H-^15^N]-TROSY spectra of ^15^N labeled dysferlin C2A samples at calcium concentrations ranging from 0 to 5 mM Ca^2+^ (Figure 3A). Instead of fast exchange (µsec) as has been reported for many C2 domains (26–28), lowering the calcium concentration pushed residues in the calcium binding loops into the immediate exchange regime. We found that lowering the calcium concentration resulted in a significant decrease in peak intensity for all three CBLs, which disappeared between 4.5 and 3 mM Ca^2+^. However, several peaks never dropped below 50% of their original value (Figure 3B). These observations suggest that these residues of dysferlin C2A may comprise a stable core region that accounts for approximately half of the protein opposite of the calcium binding loops. Interestingly, the stable core does not include all the β-strands. The ends of the strands located near the calcium binding loops show reduced intensity at lower calcium levels, indicating that these residues are also undergoing increased dynamics. Thus, the protein may be sampling an alternative state in the absence of calcium.

**Fig. 3.**
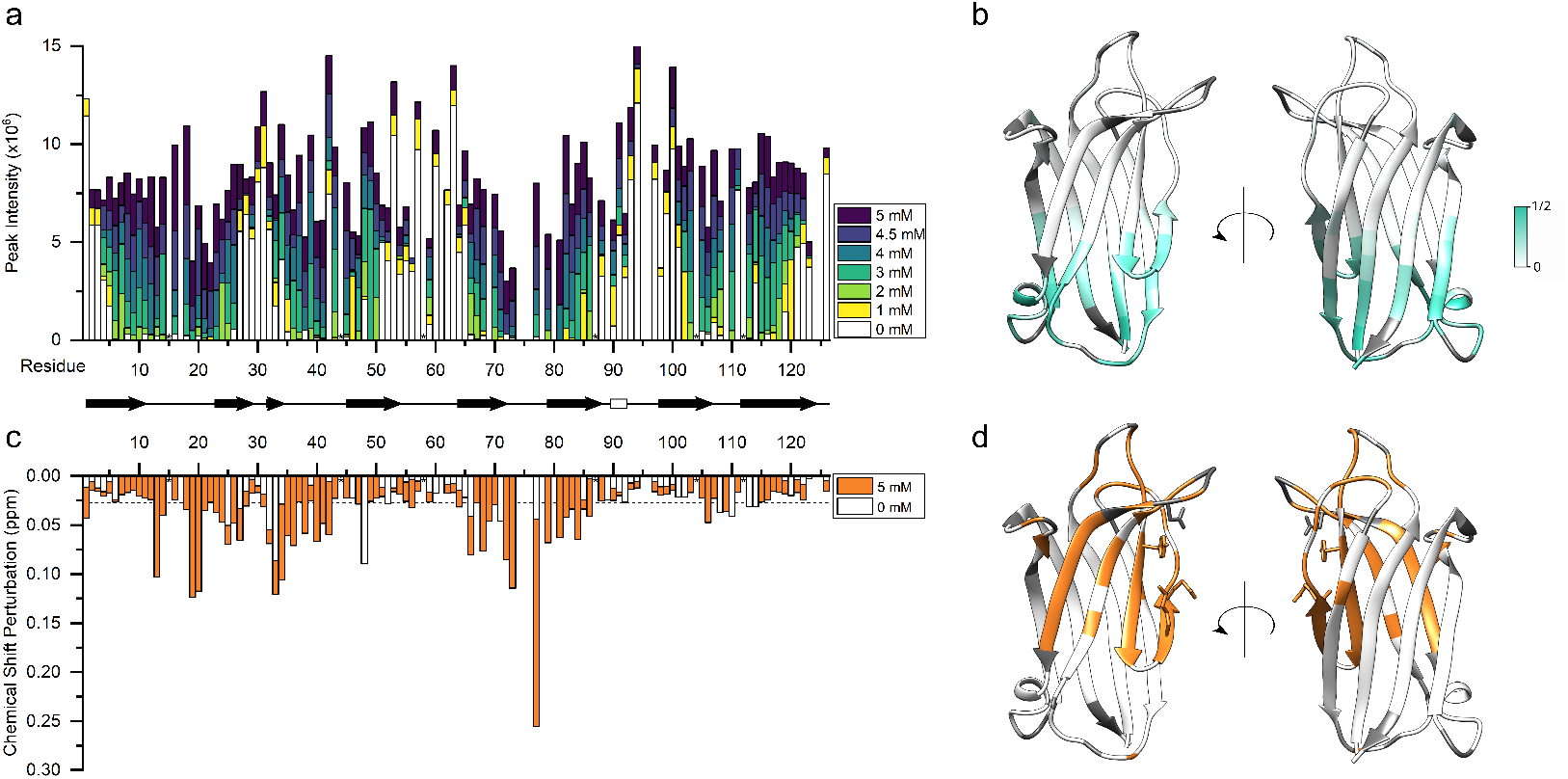
a) [^1^H-^15^N]-TROSY amide peak intensities as a function of primary sequence position in the presence of 5, 4.5, 4, 3, 2, 1, and 0 mM CaCl_2_. Proline residues indicated with an asterisk b) Ratio of 5 mM CalCl_2_ peak height to 0 mM CaCl_2_ peak height mapped onto the structure (4IHB) c) Final chemical shift perturbation as a function of primary sequence position after titration with IP3 in the presence (5 mM CaCl_2_) and absence (0.5 mM EDTA) of calcium. Proline residues indicated with an asterisk d) Significantly shifted residues mapped onto the structure (4IHB).

To further investigate the dynamic behavior of dysferlin C2A we turned to nuclear spin-relaxation measurements as a function of calcium concentration. ^15^N T_1_, T_2_ and heteronuclear NOE (HetNOE) were collected in the presence of 5, 3, and 1 mM CaCl_2_. We analyzed the spin-relaxation data using modelfree formalism (29) and the results are summarized in Figure <u>4</u>and Tables S1-3. At 5 mM CaCl_2_, the order parameters (S^2^s) are generally between 0.8 and 1.0 in regions of secondary structure, with some deviations below 0.8 in loop regions and the termini. The S^2^s decrease with decreasing calcium concentration across the protein, consistent with an overall reduction of rigidity in the structure. Samples with 1 mM calcium were found to have the lowest average S^2^ values of 0.869 ± 0.127, while 3 and 5mM calcium concentrations resulted in higher average S^2^ values of 0.897 ± 0.071 and 0.942 ± 0.061 respectively. We note that the number of residues that we could observe was reduced in the 3 and 1 mM samples as peaks began to disappear, especially in the CBL regions, indicating that these regions undergo increasing intermediate exchange as the calcium concentration decreases.

In addition to the order parameter, modelfree analysis also provides a measure of chemical exchange, R_ex_, due to motion on the millisecond to microsecond timescale. We found that the number of residues displaying an R_ex_ term and the magnitudes of the R_ex_ terms increased with decreasing calcium concentration. R_ex_ terms in the CBL regions, where measurable, increased in number and magnitude as calcium concentration decreased. We also found that several other residues showed an increase in the number and magnitude of R_ex_ terms with decreasing calcium concentration. These residues mapped to the concave face of the β-sandwich fold. Overall, these results indicate that dysferlin C2A is flexible at “low” calcium concentrations and the flexibility is progressively reduced with increasing calcium (Figure 4).

**Fig. 4.**
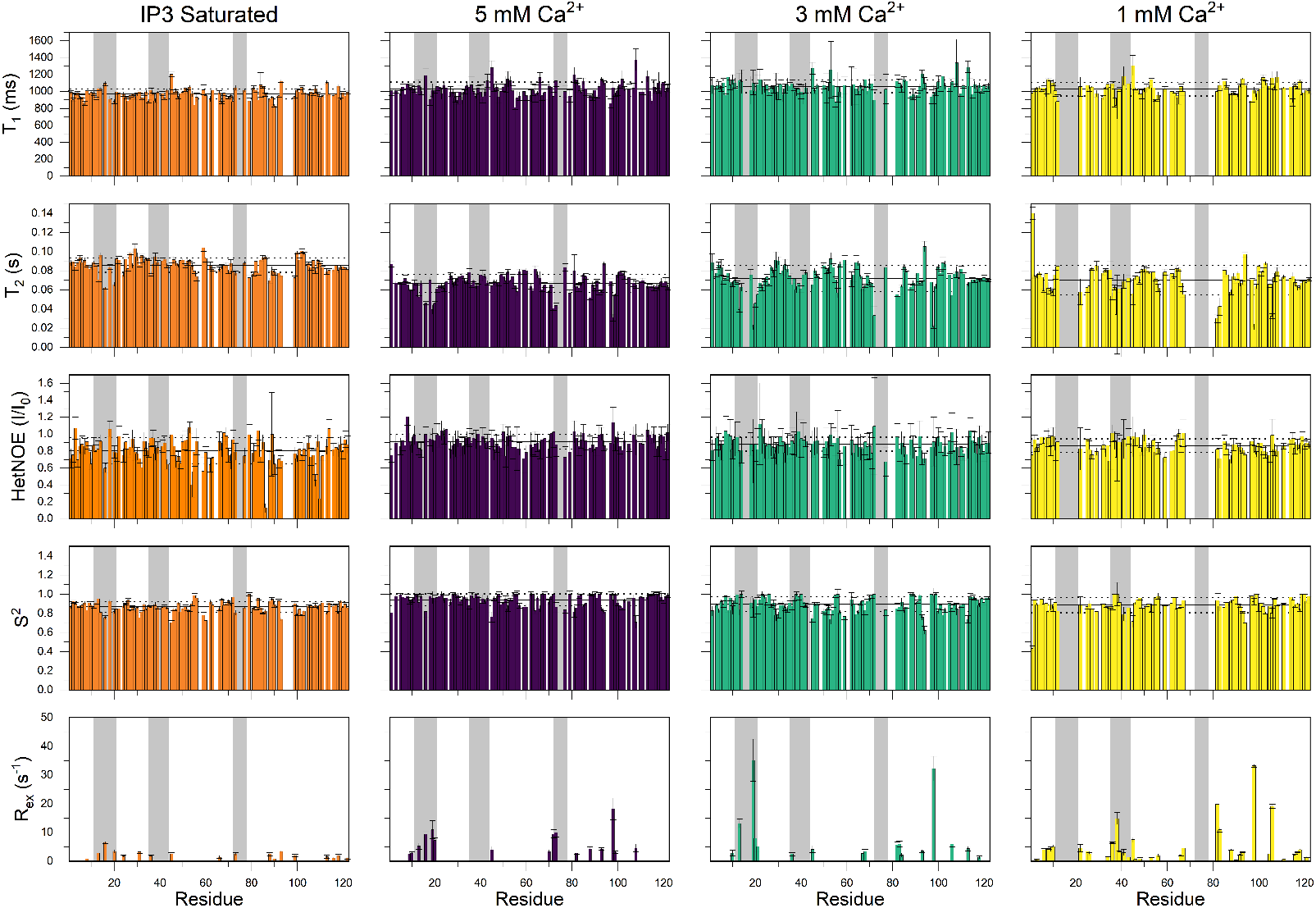
Nuclear spin-relaxation analysis for dysferlin C2A under IP3-saturating conditions, and in the presence of 5 mM, 3 mM, and 1 mM CaCl_2_. T_1_, T_2_, and HetNOE values were collected at 303K and at 800 MHz. FAST-modelfree was used to extract S^2^ and R_ex_ values. Grey vertical boxes delineate the calcium binding loops. Average values ± SD are indicated by horizontal solid and dotted lines, respectively.

### Calcium Shifts the Dynamic Equilibrium of C2A Between Two Structural States

Given our observation of calcium dependent R_ex_ terms that indicate chemical exchange processes on the microsecond to millisecond timescale, we performed chemical exchange saturation transfer (CEST) experiments to quantify these motions. ^15^N CEST experiments were performed at 1, 3, and 5 mM calcium (Figure 5 and Table 1). Chemical exchange was observed primarily in or near the CBLs at 5 mM calcium, consistent with the high order parameters and relatively few R_ex_ terms identified in our modelfree analysis. Global modeling for these residues showed that the 99.8% of the protein is in the calcium bound state, with only 0.02% in the alternate state.

**Table 1.**
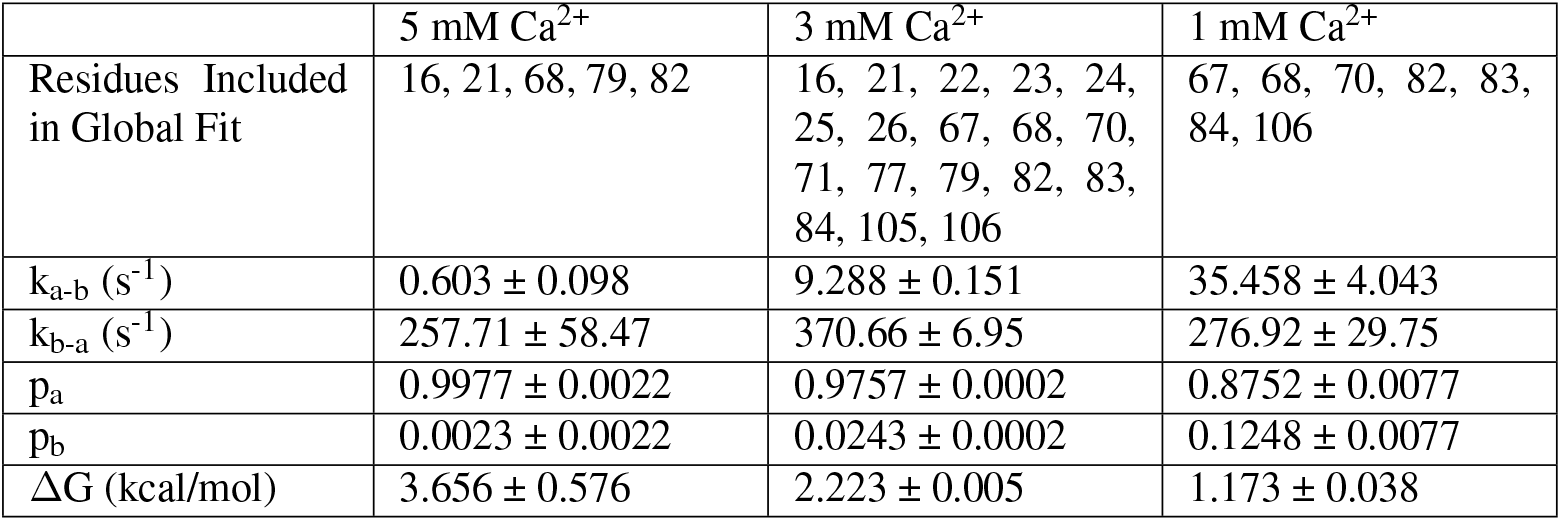
Compiled CEST-derived data from global fits.

**Fig. 5.**
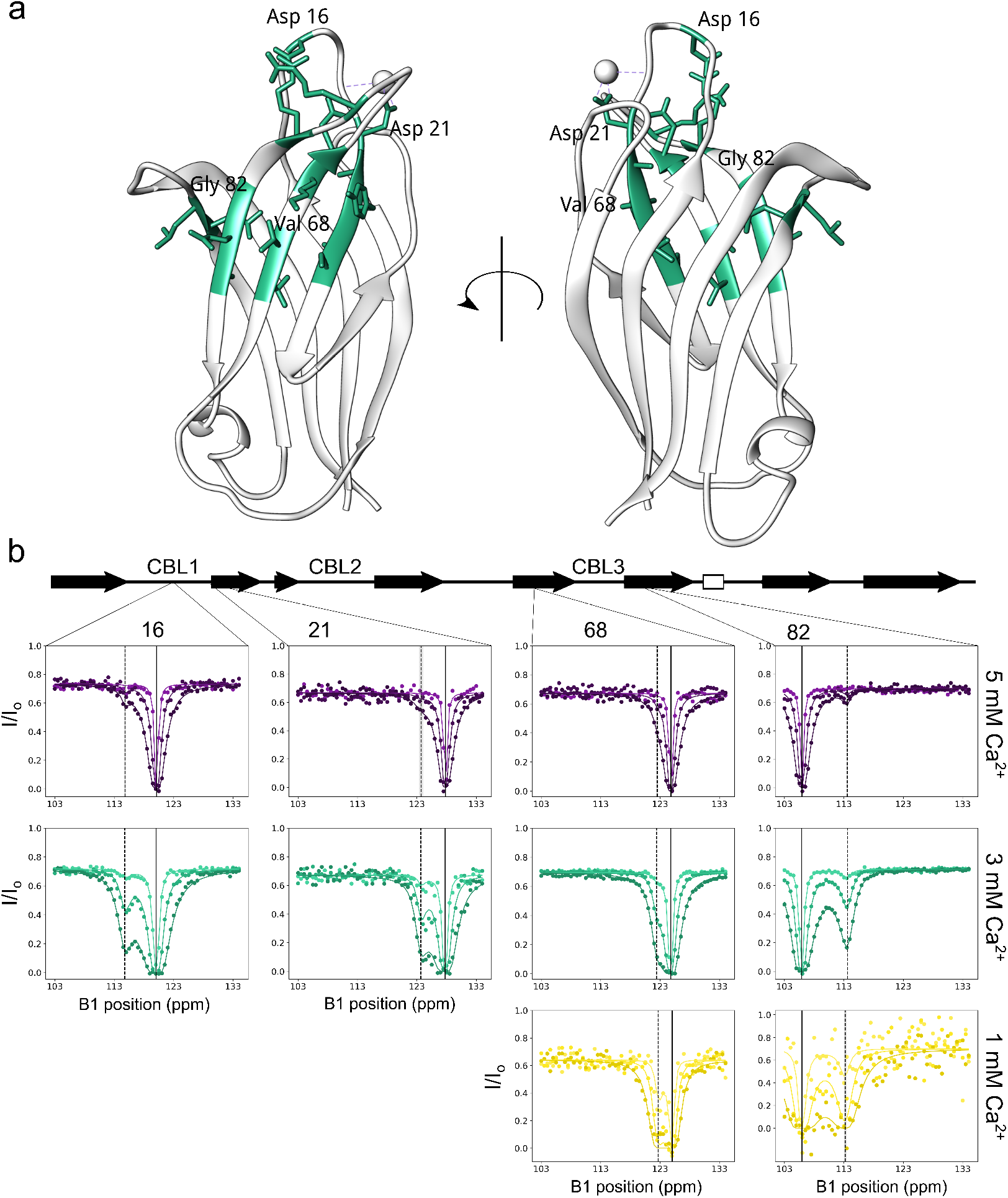
a) Residues exchanging at 3mM CaCl_2_ mapped onto the structure (4IHB). b) Selected CEST traces showing increased exchange at lower calcium concentrations. Data (circles) and fits (lines) for 50 (dark), 25 (medium), and 10 (light) Hz saturation field strengths. Solid vertical lines indicate the ^15^N chemical shift for the major state, dotted vertical lines indicate the ^15^N chemical shift for the minor state, with error indicated by grey shading.

At 3 mM calcium we found 13 residues that showed CEST profiles with chemical exchange. Each of these profiles was individually fit to a two-state model of exchange and to a global two-state exchange model as described in the methods and summarized in Table 1. All residues undergoing chemical exchange, as directly observed by CEST, were either in the CBL regions or the concave face of the β-sandwich fold. The global modeling shows that at 3 mM calcium, dysferlin C2A is 97.6% in the calcium bound state and 2.4% in the alternate state. At 1 mM calcium, many of the resonances exhibit reduced intensity, presumably due to chemical exchange. The reduced intensity of the resonances resulted in a reduction in the number of residues that we could characterize using CEST. In this case we could identify 6 residues that showed chemical exchange by CEST. Performing a similar modeling as used for the 3 mM calcium data, we determined the individual and global exchange parameters. Again, residues undergoing exchange were found to either be in the CBL regions or the concave face of the β-sandwich fold. As expected, the population of the calcium bound (87.5%) and alternate state (12.5%) shifted towards the alternate state. Our results show that there is a dynamic equilibrium between the calcium bound and alternate states even at relatively high calcium concentrations.

### The C2A Domain of Dysferlin Binds IP3 via its Polybasic Cluster

The soluble PI(4,5)P2 headgroup analog inositol 1,4,5-trisphosphate (IP3), was titrated into a ^15^N labeled sample of dysferlin C2A at 5 mM Ca^2+^. Chemical shift perturbations (CSPs) were measured over the course of the titration and final values were plotted against the residue number to map the IP3 binding site. The titration of the C2A domain with IP3 resulted in significant chemical shift perturbations, ranging from 0.255 to the 3*σ* cut-off of 0.0275 ppm, indicative of direct interaction between IP3 and C2A. Residues at the CBL3 as well as β-strands 2 and 3 demonstrated IP3 dependent CSP. This region corresponds to the concave side of the β-sandwich fold and is consistent with binding in the canonical polybasic cluster. To determine how calcium alters PIP binding to C2A, we also titrated IP3 into C2A in the absence of calcium. We found that IP3-dependent CSP’s were significantly reduced across the protein when compared to calcium-bound C2A. Of particular note, CSP’s in CBL3, which showed the largest shift upon addition of IP3 to the calcium-bound C2A, showed little or no shift without calcium present. These results show that calcium modulates the interaction surface between IP3 and C2A and is consistent with previous observations that PI(4,5)P2 binding to C2A is attenuated in the absence of calcium (13).

### The Structure of CBL3 is Stabilized by Interaction of IP3

To look into the dynamics in the IP3-bound state we collected nuclear spin-relaxation data on an IP3-saturated sample and further analyzed the data using modelfree formalism. The results are summarized in Figure <u>4</u>and Table S4. Overall, we found that the S^2^s of the IP3-saturated sample were modestly reduced. It is important to note that IP3 is known to directly interact with calcium, which could lower the free calcium concentration in the sample. Lowering the free calcium concentration would be expected to lower the S^2^ across the protein based on our dynamics data at lower calcium concentrations. In contrast to the rest of the protein, the S^2^ appears little changed or higher in CBL3 and surrounding regions in the IP3-saturated state when compared to the overall average S^2^ in the IP3-saturated state. The higher S^2^ indicates that the CBL3 is at least as rigid as the whole protein, when IP3 is bound. In contrast, in the IP3-unbound state, the S^2^ for CBL3 appears lower than the average for the rest of the protein indicating that CBL3 is less rigid than the whole protein in the absence of IP3. In addition, R_ex_ terms in CBL1 and 3 are attenuated in the IP3 saturated sample, indicating that chemical exchange at CBLs 1 and 3 is also reduced by IP3 binding, further supporting an increase in rigidity in the CBL’s upon IP3 binding. These data provide evidence that the structure of CBL3 is stabilized by interaction with IP3. Interestingly, the S^2^s for CBL2 and CBL1 do not appear to indicate an increase in rigidity in the IP3 bound state, suggesting less change in the rigidity of CBL2 and CBL1 upon IP3 binding. We note that R_ex_ terms in CBL1 are attenuated, indicating that slower, chemical exchange motions are attenuated in these regions upon IP3 binding. Overall, these data indicate that CBL3 is more directly involved in IP3 binding when compared to CBL2 and CBL1.

## Discussion

The calcium and membrane binding properties of different C2 domains vary widely, with the properties of the domain tuned to the specific physiological role of the protein. The dysferlin C2A domain differs from the equivalent domain in other ferlins in that it binds both PI(4,5)P2 and calcium. Truncation or missense mutations of the dysferlin C2A domain result in loss of dysferlin activity and the emergence of pathology, highlighting the importance of the domain. Using solution state NMR spectroscopy, we provide a structural basis for the allosteric connection between calcium and PI(4,5)P2 binding activities of the C2A domain. First we sought to characterize the domain’s response to calcium. Overall, we found that C2A has a core that is insensitive to calcium. By contrast, increasing calcium restricted the dynamics of the calcium binding loop regions and neighboring parts of the β-sandwich fold. Motion of the loops on the milito microsecond timescale was found to be calcium sensitive, as evidenced by the loss of resonance intensity in and around the calcium binding loops with decreasing calcium concentration. Interestingly, the loss of intensity extends into the β-strands proximal to the calcium binding loops, suggesting that the β-strands are also stabilized by calcium binding. This observation is further supported by the appearance of R_ex_ terms for residues in the β-strands at low calcium concentrations, indicating that residues in the β-strands are also undergoing chemical exchange. Calcium induced changes in modelfree R_ex_ values were also observed for CBL1, CBL2, and CBL3 supporting a role for calcium in affecting loop dynamics. Together, these changes indicate that coordination of calcium ions restricts the dynamics of both the apical loops and top regions of the β-strands. This change in the internal motion of the domain may favor specific structural conformations that are a prerequisite for the formation of a membrane-C2A complex, and thus, contribute to the calcium sensitivity of C2A-membrane binding.

PI(4,5)P2 binding by C2A is thought to be integral to the formation of the complex between C2A and the phospholipid membrane. Our results show that the headgroup IP3 interacts with the concave face of the β-sandwich fold and CBL3. This binding region agrees with the consensus site seen in type I C2 domains (8–11), though dysferlin C2A is of a type II fold. Tyr23 in β-strand 2, Lys32 and Arg34 in β-strand 3, and Asn78 on CBL3 are all conserved, the exception is a serine in place of a lysine at position 25. In addition, measurements of chemical exchange values revealed IP3 sensitivity within the CBL3, suggesting that a structural rearrangement of the loop may occur to accommodate IP3. Loop rearrangement would also distinguish the dysferlin C2A domain from rabphilin 3A C2A, which remains structurally unchanged when bound to PI(4,5)P2 (9).

Previous studies of C2A have shown that calcium binding enhances PI(4,5)P2 binding (13). Our results offer a molecular scale mechanism for this observation. Calcium binding to C2A stabilizes both the calcium binding loops and the β-strands near these binding loops, including Tyr23 and Asn78 which comprise the top region of the PI(4,5)P2 site. Our CEST results show that several key residues in this site undergo increasing chemical exchange as the calcium concentration decreases. These residues are clearly sampling a structurally different state, as indicated by the significant differences in chemical shifts for the excited state. However, the observed chemical shifts for the excited state are not consistent with the expected chemical shifts for an unfolded state (Figure S1). Thus, we propose that calcium binding enhances PI(4,5)P2 binding by switching C2A from a more dynamic, PI(4,5)P2 low affinity state, to a more rigid, PI(4,5)P2 high affinity state, where the β-strands in the concave face of the β-sandwich are stabilized for PI(4,5)P2 binding. Previously it was postulated that in the case of PKCα-C2, calcium binding enhanced PI(4,5)P2 binding by facilitating access to the polybasic cluster (30). Our findings provide support for a different mechanism: the low calcium concentrations in un-stimulated cells would allow the C2A domain to occupy a dynamic, low PI(4,5)P2 affinity state and thus would be predominantly unbound from membranes. Upon calcium influx, through microtears in the sarcolemma or through DHPR or ryanodine receptors in the transverse tubules, C2A is stabilized and converted from the low PI(4,5)P2 affinity state to the high PI(4,5)P2 affinity state, resulting in membrane binding and activation of dysferlin (Figure 6b).

**Fig. 6.**
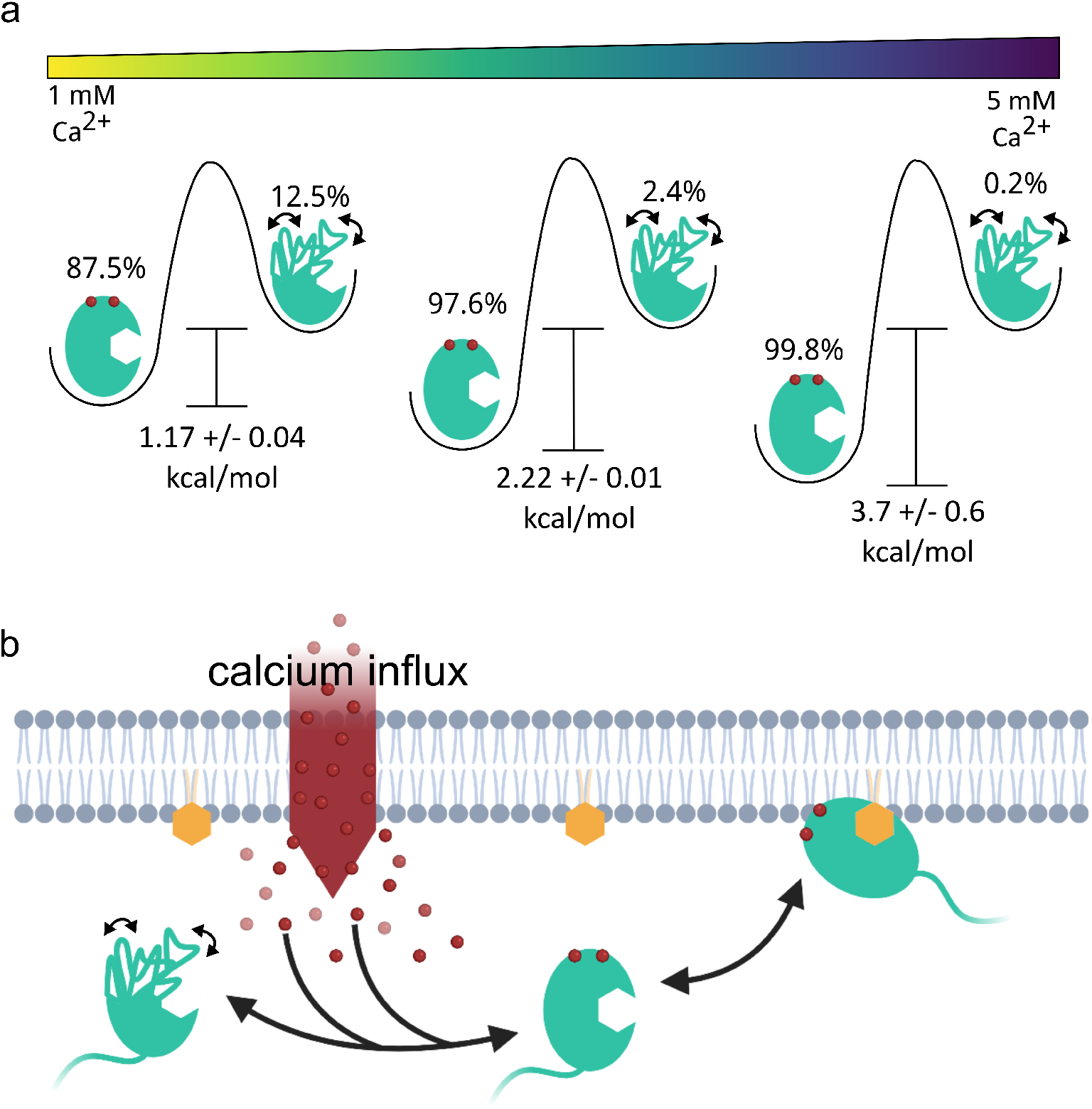
Proposed model for dysferlin C2A action. a) Increasing calcium concentrations stabilize the calcium binding loops and adjacent regions of the domain, shifting the equilibrium to a conformation that favors PI(4,5)P2 binding b) Within the cell, calcium can trigger the C2A conformational change, which subsequently results in dysferlin anchoring to the membrane. Created with BioRender.com.

Based on our measurements, we propose that the C2A docks with the membrane at an acute angle that includes contact between β-strands 2 and 3 on the domain and the PI(4,5)P2 headgroup in the membrane. A reorientation of CBL3 appears to help facilitate binding with the lipid headgroup, however this does not inhibit calcium ion coordination by the domain. The calcium binding region of many C2 domains stabilizes domain interaction with phosphatidylserine through a calcium bridging mechanism, and liposome binding assays support a similar bridging effect for dysferlin C2A (6). The inositol phosphate sensitive residues identified in our study are distinct from the region of the domain that interacts with phosphatidylserine, allowing for simultaneous membrane interactions in the presence of calcium. The observed reduction in dynamics of the CBL region of the domain upon calcium binding may serve to preorganize the loops for interaction with the phosphatidylserine lipid, and thus serve as a second method for conferring calcium sensitivity to membrane binding.

Dysferlin is a broadly conserved membrane trafficking protein, present from sea stars (31) to humans (32, 33). Mutations in dysferlin have been directly linked to muscular dystrophies and cardiomyopathy, and dysferlin-null mouse models display muscular dystrophy-like pathologies (34). The large size of the full-length protein prohibits its utility in gene delivery therapies (35), thus we must be judicious in our design of smaller constructs more appropriate for gene delivery. A truncated dysferlin composed of the only the two most C-terminal C2 domains can rescue the membrane resealing deficiency, but it is not sufficient to completely alleviate the dystrophic phenotype in mice (36). This indicates that there is perhaps an equally important role for the N-terminal dysferlin domains in muscle tissue. Here our work argues for the importance of the N-terminal C2A domain and provides a rationale for including the domain in future gene therapy constructs.

It remains to be seen how mutations, such as V67D, that lie far from the calcium binding loops and the canonical polybasic cluster, impact the two states, however the calcium sensitive exchange beyond these regions we have identified may provide the explanation for the resulting pathology. Furthermore, while we have shown how calcium binding modulates PI(4,5)P2 affinity, interaction with other C2A binding partners may also influence the populations of the exchanging states, leading to modulation of activity. Further refinement of structural methods for probing protein-membrane interaction, like those demonstrated in this study, may shed light on these future directions.

## Materials and Methods

### Molecular Biology and Protein Purification

Human dysferlin cDNA (AF075575), a gift from Dr. Kate Bushby (Newcastle University), was used as a template for cloning. The C2A domain (amino acids 1-129) was cloned into the pET-28a(+) vector (Novagen) between the BamHI and HindIII restriction sites. Complementary flanking restriction sites for the dysferlin insert were generated by PCR amplification using the following primers: forward 5’-GCG CGC GGA TCC ATG CTG AGG GTC TTC ATC CTC TAT GCC-3’ reverse 5’-GCG CGC AAG CTT TTA AGC TCC AGG CAG CGG-3’. BL21 DE3 cells harboring the expression plasmid were cultured overnight at 37°C in Luria-Bertani broth containing 50 µg/mL kanamycin and 1% w/v glucose and were used to seed 1 liter cultures of Luria-Bertaini broth containing 50 µg/mL kanamycin at a ratio of 1:1000. ^15^N and ^15^N/^13^C isotopically labeled protein was produced by bacteria grown in MJ9 media containing 1 g/L ^15^NH_4_Cl and 2 g/L of ^12^C or ^13^C glucose. These were then grown to an optical density of 0.6 at 37°C and induced with 0.5 mM isopropyl β-D-1-thiogalactopyranoside (IPTG). Protein was expressed for 16 hours at 18°C. Cultures were centrifuged at 4000 rpm at 4°C for 20 minutes and resuspended in lysis buffer: 50 mM HEPES pH 8, 250 mM NaCl, 10% (v/v) glycerol, 5 mM CaCl_2_, 1 mM phenylmethanesulfonyl fluoride (PMSF), and 1 µM leupeptin, pepstatin A, and aprotonin. Cells were lysed using a Microfluidics M-11P microfluidizer at 18,000 psi. 0.5% CHAPS (w/v) was added to the total lysate and left to rock for 1 hour on ice. Soluble fractions were then obtained by centrifugation in a Beckman J2-21 centrifuge at 20,000 x g at 4°C for 20 minutes. Clarified lysate was bound to His-Pur Cobalt resin (Thermo Scientific) for 2 hours with rocking at 4°C. Beads were then washed with 20 column volumes of lysis buffer and protein was eluted with 50 mM HEPES, 150 mM NaCl, 5 mM CaCl_2_, and 200 mM imidazole. Purity of the elution fractions was confirmed via SDS-PAGE, pooled, and dialyzed against 50 mM HEPES, 150 mM NaCl, 5 mM CaCl_2_ and 1 mM 1,4-dithiothreitol (DTT). The hexahistidine purification tag was cleaved from isotopically labeled samples by thrombin during dialysis into a low salt buffer (50 mM HEPES, 20 mM NaCl, 5 mM CaCl_2_, 1 mM DTT, pH 7). The cleaved protein was further purified by cation exchange chromatography (HiTrap(tm) SP HP, GE). Samples were then dialyzed into NMR buffer (50 mM HEPES, 100 mM KCl, 5 mM CaCl_2_, 5 mM DTT, pH 7), concentrated, and supplemented with 1 mM sodium azide, 10% D_2_O, and 2-2 dimethylsilapentane-5-sulfonic acid prior to data collection.

### NMR Backbone Assignments

NMR spectra were collected at 303K on a 800-MHz Bruker Avance IIIHD spectrometer equipped with a TCI cryoprobe. Backbone resonance assignments were determined from a set of [^15^N, ^1^H] TROSY based triple-resonance experiments. (HNCA, HNCACB, HNCOCACB, HNCO, HNCACO). NMR spectra were processed with NMRPipe (37) and analyzed with CcpNmr Analysis (38). TALOS+ was used to calculate secondary structure propensity (39). Chemical shifts have been deposited with the BMRB under accession number 50753.

### NMR Titrations

Peak intensities for the titration experiments were normalized to the DSS signal to correct for changes in sample concentration. For the calcium titration a sample in 5 mM CaCl_2_ was back titrated with 0.5 mM EDTA (ethylenediaminetetraacetic acid) dissolved in NMR buffer lacking CaCl_2_. D-myo-Inositol 1,4,5-tris-phosphate trisodium salt (Sigma-Aldrich) was dissolved in appropriate NMR buffer to stock concentrations of 2.057 or 20.57 mM and titrated to a final IP3:C2A ratio of 7.6:1.

### NMR Spin Relaxation

T_1_ and T_2_ ^15^N relaxation times and steady-state ^15^N-^1^H NOE values for all samples were obtained using standard [^15^N, ^1^H] TROSY based Bruker pulse sequences. (trt1etf3gpsitc3d, trt2eft3gpsitc3d, trnoef3gpsi) Relaxation delay times for T_1_ experiments varied from 20 ms to 1.2 s, relaxation delay times for T_2_ experiments ranged from 33.9 ms to 271.4 ms. The steady-state ^15^N-H NOE experiment included a total recovery delay of 8 seconds. Peak intensities were measured as peak heights, which were fit to a single exponential, I=I_0_e^(-rate*t)^, using CcpNMR Analysis to extract R_1_ and R_2_ values. NOE values were calculated by dividing peak intensity in the presence of amide saturation by peak intensity in the absence of amide saturation. Error (σ) in NOE values was calculated using the equation σ/NOE = [(δ_unsat_/I_unsat_)^2^+(δ_sat_/I_sat_)^2^] where δ is the baseline noise in each spectrum. Quadric diffusion (40) was used to determine the diffusion tensor with PDB model 4IHB. We used FAST-Modelfree to extract generalized order parameters and R_ex_ terms from spin-relaxation data (29, 41)

### Chemical Exchange Saturation Transfer (CEST) Measurements

^15^N CEST data was collected using a standard Bruker pulse sequence (hsqc_cest_etf3gpsitc3d) with saturation field strengths of 50, 25, and 10 hz, with a saturation time of 0.4 seconds, and a 1.5 second recovery delay. A total of 64 planes were collected, 0.5 ppm steps from 0 ppm and 134.0 ppm and one reference plane. Fits to CEST intensity profiles were done using the ChemEx program (https://github.com/gbouvignies/chemex) (42). Individual fits were made for each residue to a two-state exchange model, then residues where the error in exchange rate exceeded the exchange rate itself were rejected. The remaining residues yielded similar populations and exchange rates. We subsequently globally fit these residues to the two-state exchange model, yielding global exchange parameters (Table 1). Gibbs free energies were calculated from the population differences. To identify additional residues that were in exchange, but where the chemical shift differences were small, we further fit the data for residues close to those previously fit in the global model, using the global exchange rate and populations as fixed parameters. This analysis resulted in the identification of 4 additional exchanging residues in the 3 mM Ca^2+^ sample.

## ACKNOWLEDGEMENTS

We thank Elisar Barbar for constructive feedback. This work was supported by the NIH National Institute of Deafness and Other Communication Disorders (NIDCD) Grant R01DC014588 and the National Science Foundation award CHE 1905091 and MCB 2019386. The Oregon State University NMR Facility is supported in part by the National Institutes of Health, HEI Grant 1S10OD018518 and by M. J. Murdock Charitable Trust Grant 2014162.

## Bibliography

1. Eric A. Nalefski and Joseph J. Falke. The C2 domain calcium-binding motif: Structural and functional diversity. Protein Science, 5(12):2375–2390, December 1996. ISSN 09618368, 1469896X. doi: 10.1002/pro.5560051201.

2. Josep Rizo and Thomas C. Südhof. C 2 -domains, Structure and Function of a Universal Ca 2+ -binding Domain. Journal of Biological Chemistry, 273(26):15879–15882, June 1998. ISSN 0021-9258, 1083-351X. doi: 10.1074/jbc.273.26.15879.

3. X. Shao, I. Fernandez, T. C. Südhof, and J. Rizo. Solution structures of the Ca2+-free and Ca2+-bound C2A domain of synaptotagmin I: does Ca2+ induce a conformational change? Biochemistry, 37(46):16106–16115, November 1998. ISSN 0006-2960. doi: 10.1021/bi981789h.

4. Josep Ubach, Xiangyang Zhang, Xuguang Shao, Thomas C. Südhof, and Josep Rizo. Ca2+ binding to synaptotagmin: how many Ca2+ ions bind to the tip of a C2-domain? The EMBO Journal, 17(14):3921–3930, July 1998. ISSN 0261-4189, 1460-2075. doi: 10.1093/emboj/17.14.3921.

5. Naomi J. Marty, Chelsea L. Holman, Nazish Abdullah, and Colin P. Johnson. The C2 Domains of Otoferlin, Dysferlin, and Myoferlin Alter the Packing of Lipid Bilayers. Biochemistry, 52(33):5585–5592, August 2013. ISSN 0006-2960. doi: 10.1021/bi400432f. Publisher: American Chemical Society.

6. Nazish Abdullah, Murugesh Padmanarayana, Naomi J. Marty, and Colin P. Johnson. Quantitation of the Calcium and Membrane Binding Properties of the C2 Domains of Dysferlin. Biophysical Journal, 106(2):382–389, January 2014. ISSN 0006-3495. doi: 10.1016/j.bpj.2013.11.4492. Publisher: Elsevier.

7. W Cho and R Stahelin. Membrane binding and subcellular targeting of C2 domains. Biochimica et Biophysica Acta (BBA) - Molecular and Cell Biology of Lipids, 1761(8):838– 849, August 2006. ISSN 13881981. doi: 10.1016/j.bbalip.2006.06.014.

8. Marta Guerrero-Valero, Cristina Ferrer-Orta, Jordi Querol-Audí, Consuelo Marin-Vicente, Ignacio Fita, Juan C. Gómez-Fernández, Nuria Verdaguer, and Senena Corbalán-García. Structural and mechanistic insights into the association of PKCα-C2 domain to PtdIns(4,5)P2. Proceedings of the National Academy of Sciences, 106(16):6603–6607, April 2009. ISSN 0027-8424, 1091-6490. doi: 10.1073/pnas.0813099106. Publisher: National Academy of Sciences Section: Biological Sciences.

9. Jaime Guillén, Cristina Ferrer-Orta, Mònica Buxaderas, Dolores Pérez-Sánchez, Marta Guerrero-Valero, Ginés Luengo-Gil, Joan Pous, Pablo Guerra, Juan C. Gómez-Fernández, Nuria Verdaguer, and Senena Corbalán-García. Structural insights into the Ca2+ and PI(4,5)P2 binding modes of the C2 domains of rabphilin 3A and synaptotagmin 1. Proceedings of the National Academy of Sciences, 110(51):20503–20508, December 2013. ISSN 0027-8424, 1091-6490. doi: 10.1073/pnas.1316179110. Publisher: National Academy of Sciences Section: Biological Sciences.

10. Chanjuan Wan, Bo Wu, Zhenwei Song, Jiahai Zhang, Huiying Chu, Aoli Wang, Qingsong Liu, Yunyu Shi, Guohui Li, and Junfeng Wang. Insights into the molecular recognition of the granuphilin C2A domain with PI(4,5)P2. Chemistry and Physics of Lipids, 186:61–67, February 2015. ISSN 0009-3084. doi: 10.1016/j.chemphyslip.2015.01.003.

11. Lirin Michaeli, Irit Gottfried, Maria Bykhovskaia, and Uri Ashery. Phosphatidylinositol (4, 5)-bisphosphate targets double C2 domain protein B to the plasma membrane. Traffic (Copenhagen, Denmark), 18(12):825–839, December 2017. ISSN 1600-0854. doi: 10.1111/tra.12528.

12. Sonia Sánchez-Bautista, Consuelo Marín-Vicente, Juan C. Gómez-Fernández, and Senena Corbalán-García. The C2 domain of PKCalpha is a Ca2+ -dependent PtdIns(4,5)P2 sensing domain: a new insight into an old pathway. Journal of Molecular Biology, 362(5):901–914, October 2006. ISSN 0022-2836. doi: 10.1016/j.jmb.2006.07.093.

13. Christian Therrien, Sabrina Di Fulvio, Sarah Pickles, and Michael Sinnreich. Characterization of Lipid Binding Specificities of Dysferlin C2 Domains Reveals Novel Interactions with Phosphoinositides †. Biochemistry, 48(11):2377–2384, March 2009. ISSN 0006-2960, 1520-4995. doi: 10.1021/bi802242r.

14. Jihong Bai, Ward C. Tucker, and Edwin R. Chapman. PIP 2 increases the speed of response of synaptotagmin and steers its membrane-penetration activity toward the plasma membrane. Nature Structural & Molecular Biology, 11(1):36–44, January 2004. ISSN 1545-9985. doi: 10.1038/nsmb709. Number: 1 Publisher: Nature Publishing Group.

15. Chie Matsuda, Katsuya Miyake, Kimihiko Kameyama, Etsuko Keduka, Hiroshi Takeshima, Toru Imamura, Nobukazu Araki, Ichizo Nishino, and Yukiko Hayashi. The C2A domain in dysferlin is important for association with MG53 (TRIM72). PLOS Currents Muscular Dystrophy, November 2012. ISSN 2157-3999. doi: 10.1371/5035add8caff4. Publisher: Public Library of Science.

16. W.-Q. Han, M. Xia, M. Xu, K. M. Boini, J. K. Ritter, N.-J. Li, and P.-L. Li. Lysosome fusion to the cell membrane is mediated by the dysferlin C2A domain in coronary arterial endothelial cells. Journal of Cell Science, 125(5):1225–1234, March 2012. ISSN 0021-9533, 1477-9137. doi: 10.1242/jcs.094565.

17. Julia Hofhuis, Kristina Bersch, Ronja Büssenschütt, Marzena Drzymalski, David Liebetanz, Viacheslav O. Nikolaev, Stefan Wagner, Lars S. Maier, Jutta Gärtner, Lars Klinge, and Sven Thoms. Dysferlin mediates membrane tubulation and links T-tubule biogenesis to muscular dystrophy. Journal of Cell Science, 130(5):841–852, March 2017. ISSN 0021-9533, 1477-9137. doi: 10.1242/jcs.198861. Publisher: The Company of Biologists Ltd Section: Research Article.

18. Joseph A. Roche, Lisa W. Ru, Andrea M. O’Neill, Wendy G. Resneck, Richard M. Lovering, and Robert J. Bloch. Unmasking Potential Intracellular Roles For Dysferlin through Improved Immunolabeling Methods. Journal of Histochemistry & Cytochemistry, 59(11):964–975, November 2011. ISSN 0022-1554, 1551-5044. doi: 10.1369/0022155411423274.

19. C. Hidalgo, J. Jorquera, V. Tapia, and P. Donoso. Triads and transverse tubules isolated from skeletal muscle contain high levels of inositol 1,4,5-trisphosphate. Journal of Biological Chemistry, 268(20):15111–15117, July 1993. ISSN 0021-9258. doi: 10.1016/S0021-9258(18)82444-9.

20. Beryl N. Ampong, Michihiro Imamura, Teruhiro Matsumiya, Mikiharu Yoshida, and Shin’ichi Takeda. Intracellular localization of dysferlin and its association with the dihydropyridine receptor. Acta Myologica: Myopathies and Cardiomyopathies: Official Journal of the Mediterranean Society of Myology, 24(2):134–144, October 2005. ISSN 1128-2460.

21. Jaclyn P. Kerr, Christopher W. Ward, and Robert J. Bloch. Dysferlin at transverse tubules regulates Ca2+ homeostasis in skeletal muscle. Frontiers in Physiology, 5, March 2014. ISSN 1664-042X. doi: 10.3389/fphys.2014.00089.

22. Sandra T. Cooper and Paul L. McNeil. Membrane Repair: Mechanisms and Pathophysiology. Physiological Reviews, 95(4):1205–1240, October 2015. ISSN 0031-9333, 1522-1210. doi: 10.1152/physrev.00037.2014.

23. Alexis R. Demonbreun, Madison V. Allen, James L. Warner, David Y. Barefield, Swathi Krishnan, Kaitlin E. Swanson, Judy U. Earley, and Elizabeth M. McNally. Enhanced Muscular Dystrophy from Loss of Dysferlin Is Accompanied by Impaired Annexin A6 Translocation after Sarcolemmal Disruption. The American Journal of Pathology, 186(6):1610–1622, June 2016. ISSN 1525-2191. doi: 10.1016/j.ajpath.2016.02.005.

24. Kerry Fuson, Anne Rice, Ryan Mahling, Adam Snow, Kamakshi Nayak, Prajna Shanbhogue, Austin G. Meyer, Gregory M.I. Redpath, Anne Hinderliter, Sandra T. Cooper, and R. Bryan Sutton. Alternate Splicing of Dysferlin C2A Confers Ca2+-Dependent and Ca2+-Independent Binding for Membrane Repair. Structure, 22(1):104–115, January 2014. ISSN 09692126. doi: 10.1016/j.str.2013.10.001.

25. Yuning Wang, Roya Tadayon, Liliana Santamaria, Pascal Mercier, Chantal J. Forristal, and Gary S. Shaw. Calcium binds and rigidifies the dysferlin C2A domain in a tightly coupled manner. Biochemical Journal, 478(1):197–215, January 2021. ISSN 0264-6021. doi: 10.1042/BCJ20200773.

26. Xuguang Shao, Bazbek A. Davletov, R. Bryan Sutton, Thomas C. Südhof, and Josep Rizo. Bipartite Ca2+-Binding Motif in C2 Domains of Synaptotagmin and Protein Kinase C. Science, 273(5272):248–251, July 1996. ISSN 0036-8075, 1095-9203. doi: 10.1126/science.273.5272.248. Publisher: American Association for the Advancement of Science Section: Reports.

27. Jesus Garcia, Stefan H. Gerber, Shuzo Sugita, Thomas C. Südhof, and Josep Rizo. A conformational switch in the Piccolo C2A domain regulated by alternative splicing. Nature Structural & Molecular Biology, 11(1):45–53, January 2004. ISSN 1545-9993. doi: 10.1038/nsmb707.

28. Hiromasa Yagi, Paul J. Conroy, Eleanor W. W. Leung, Ruby H. P. Law, Joseph A. Trapani, Ilia Voskoboinik, James C. Whisstock, and Raymond S. Norton.Structural Basis for Ca2+-mediated Interaction of the Perforin C2 Domain with Lipid Membranes. The Journal of Biological Chemistry, 290(42):25213–25226, October 2015. ISSN 0021-9258. doi: 10.1074/jbc.M115.668384.

29. A. M. Mandel, M. Akke, and A. G. Palmer. Backbone dynamics of Escherichia coli ribonuclease HI: correlations with structure and function in an active enzyme. Journal of Molecular Biology, 246(1):144–163, February 1995. ISSN 0022-2836. doi: 10.1006/jmbi.1994.0073.

30. Sonia Sánchez-Bautista, Consuelo Marín-Vicente, Juan C. Gómez-Fernández, and Senena Corbalán-García. The C2 Domain of PKCα Is a Ca2+-dependent PtdIns(4,5)P2 Sensing Domain: A New Insight into an Old Pathway. Journal of Molecular Biology, 362(5):901–914, October 2006. ISSN 0022-2836. doi: 10.1016/j.jmb.2006.07.093.

31. Nathalie Oulhen, Thomas M. Onorato, Isabela Ramos, and Gary M. Wessel. Dysferlin is essential for endocytosis in the sea star oocyte. Developmental biology, 388(1):94–102, April 2014. ISSN 0012-1606. doi: 10.1016/j.ydbio.2013.12.018.

32. Rumaisa Bashir, Stephen Britton, Tom Strachan, Sharon Keers, Elizabeth Vafiadaki, Majlinda Lako, Isabelle Richard, Sylvie Marchand, Nathalie Bourg, Zohar Argov, Menachem Sadeh, Ibrahim Mahjneh, Giampiero Marconi, Maria Rita Passos-Bueno, Eloisa de S Moreira, Mayana Zatz, Jacques S. Beckmann, and Kate Bushby. A gene related to Caenorhabditis elegans spermatogenesis factor fer-1 is mutated in limb-girdle muscular dystrophy type 2B. Nature Genetics, 20(1):37–42, September 1998. ISSN 1061-4036, 1546-1718. doi: 10.1038/1689.

33. Jing Liu, Masashi Aoki, Isabel Illa, Chenyan Wu, Michel Fardeau, Corrado Angelini, Carmen Serrano, J. Andoni Urtizberea, Faycal Hentati, Mongi Ben Hamida, Saeed Bohlega, Edward J. Culper, Anthony A. Amato, Karen Bossie, Joshua Oeltjen, Khemissa Bejaoui, Diane McKenna-Yasek, Betsy A. Hosler, Erwin Schurr, Kiichi Arahata, Pieter J. de Jong, and Robert H. Brown. Dysferlin, a novel skeletal muscle gene, is mutated in Miyoshi myopathy and limb girdle muscular dystrophy. Nature Genetics, 20(1):31–36, September 1998. ISSN 1061-4036, 1546-1718. doi: 10.1038/1682.

34. Reginald E. Bittner, Louise V.B. Anderson, Elke Burkhardt, Rumaisa Bashir, Elizabeth Vafiadaki, Silva Ivanova, Thomas Raffelsberger, Isabel Maerk, Harald Höger, Martin Jung, Mohsen Karbasiyan, Maria Storch, Hans Lassmann, Jennifer A. Moss, Keith Davison, Ruth Harrison, Kate M.D. Bushby, and André Reis. Dysferlin deletion in SJL mice (SJL-Dysf) defines a natural model for limb girdle muscular dystrophy 2B. Nature Genetics, 23(2): 141–142, October 1999. ISSN 1061-4036, 1546-1718. doi: 10.1038/13770.

35. Telmo Llanga, Nadia Nagy, Laura Conatser, Catherine Dial, R. Bryan Sutton, and Matthew L. Hirsch. Structure-Based Designed Nano-Dysferlin Significantly Improves Dysferlinopathy in BLA/J Mice. Molecular Therapy: The Journal of the American Society of Gene Therapy, 25(9):2150–2162, September 2017. ISSN 1525-0024. doi: 10.1016/j.ymthe.2017.05.013.

36. William Lostal, Marc Bartoli, Carinne Roudaut, Nathalie Bourg, Martin Krahn, Marina Pryadkina, Perrine Borel, Laurence Suel, Joseph A. Roche, Daniel Stockholm, Robert J. Bloch, Nicolas Levy, Rumaisa Bashir, and Isabelle Richard. Lack of Correlation between Outcomes of Membrane Repair Assay and Correction of Dystrophic Changes in Experimental Therapeutic Strategy in Dysferlinopathy. PLoS ONE, 7(5):e38036, May 2012. ISSN 1932-6203. doi: 10.1371/journal.pone.0038036.

37. Frank Delaglio, Stephan Grzesiek, Geerten W. Vuister, Guang Zhu, John Pfeifer, and Ad Bax. NMRPipe: A multidimensional spectral processing system based on UNIX pipes. Journal of Biomolecular NMR, 6(3):277–293, November 1995. ISSN 1573-5001. doi: 10.1007/BF00197809.

38. Wim F. Vranken, Wayne Boucher, Tim J. Stevens, Rasmus H. Fogh, Anne Pajon, Miguel Llinas, Eldon L. Ulrich, John L. Markley, John Ionides, and Ernest D. Laue. The CCPN data model for NMR spectroscopy: development of a software pipeline. Proteins, 59(4):687–696, June 2005. ISSN 1097-0134. doi: 10.1002/prot.20449.

39. Yang Shen, Frank Delaglio, Gabriel Cornilescu, and Ad Bax. TALOS+: a hybrid method for predicting protein backbone torsion angles from NMR chemical shifts. Journal of Biomolecular NMR, 44(4):213–223, August 2009. ISSN 1573-5001. doi: 10.1007/s10858-009-9333-z.

40. Larry K. Lee, Mark Rance, Walter J. Chazin, and Arthur G. Palmer. Rotational diffusion anisotropy of proteins from simultaneous analysis of 15N and 13Cα nuclear spin relaxation. Journal of Biomolecular NMR, 9(3):287–298, April 1997. ISSN 1573-5001. doi: 10.1023/A:1018631009583.

41. Roger Cole and J. Patrick Loria. FAST-Modelfree: A program for rapid automated analysis of solution NMR spin-relaxation data. Journal of Biomolecular NMR, 26(3):203–213, July 2003. ISSN 1573-5001. doi: 10.1023/A:1023808801134.

42. Pramodh Vallurupalli, Guillaume Bouvignies, and Lewis E. Kay. Studying “Invisible” Excited Protein States in Slow Exchange with a Major State Conformation. Journal of the American Chemical Society, 134(19):8148–8161, May 2012. ISSN 0002-7863. doi: 10.1021/ja3001419. Publisher: American Chemical Society.

