## Supplemental Figures and Tables for "Calcium Sensitive Allostery Regulates the PI(4,5)P2 Binding Site of the Dysferlin C2A Domain"

### Supplemental Materials

Contains Supplemental Figures 1-5 Supplemental Tables 1-7

DRAFT

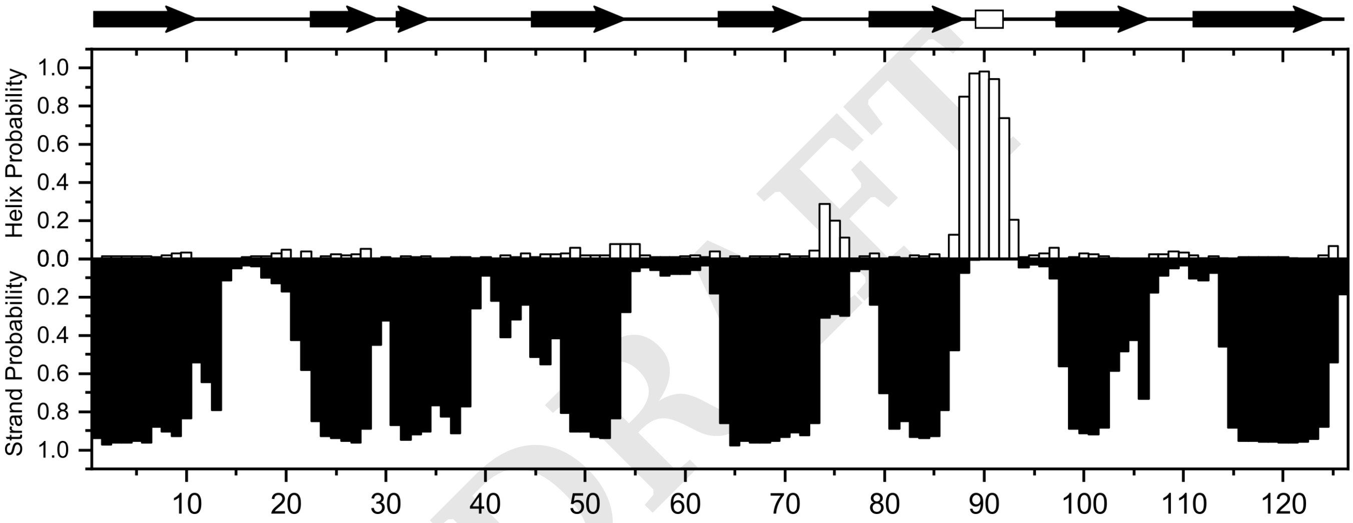

**Fig. 1.** (Top) Domain secondary structure boundaries taken from crystal structure 4IHB. (Bottom) Secondary structure propensity in solution as calculated by TALOS+.

DRAFT

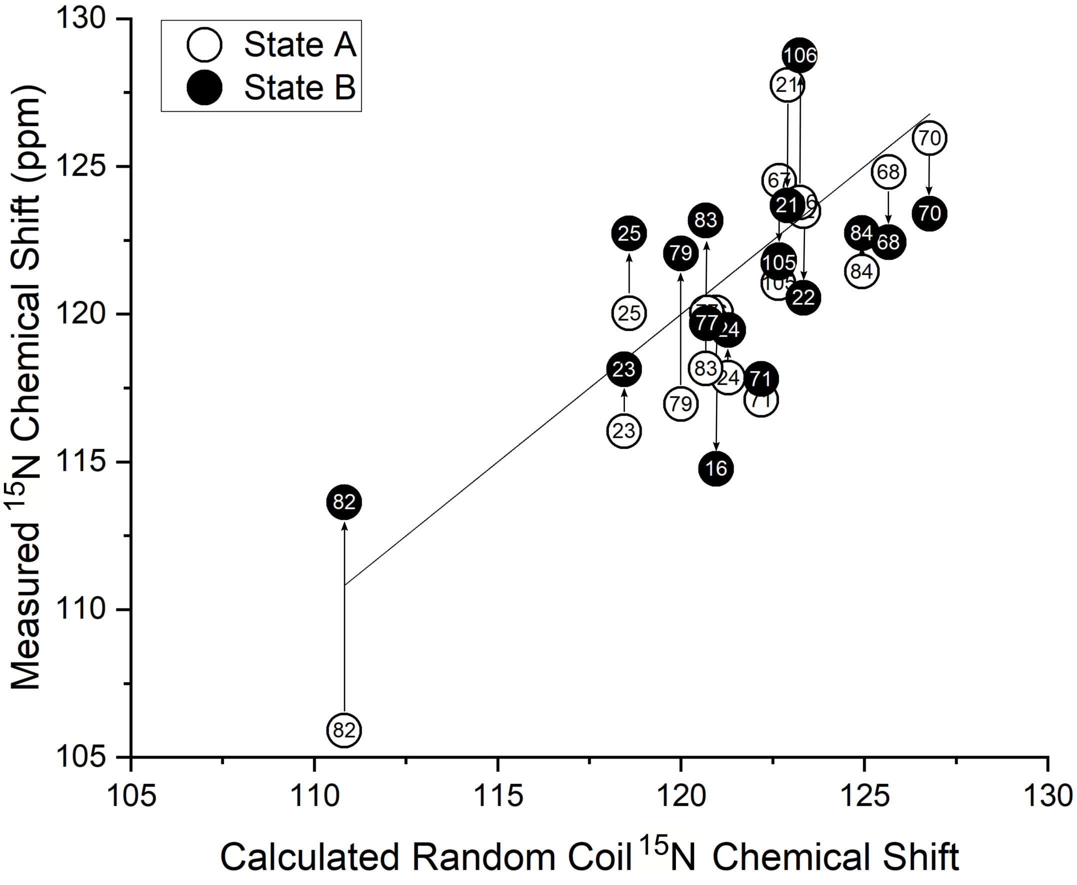

**Fig. 2.** Backbone state A and state B 15N shifts for exchanging residues Solid line is 1:1 correlation.

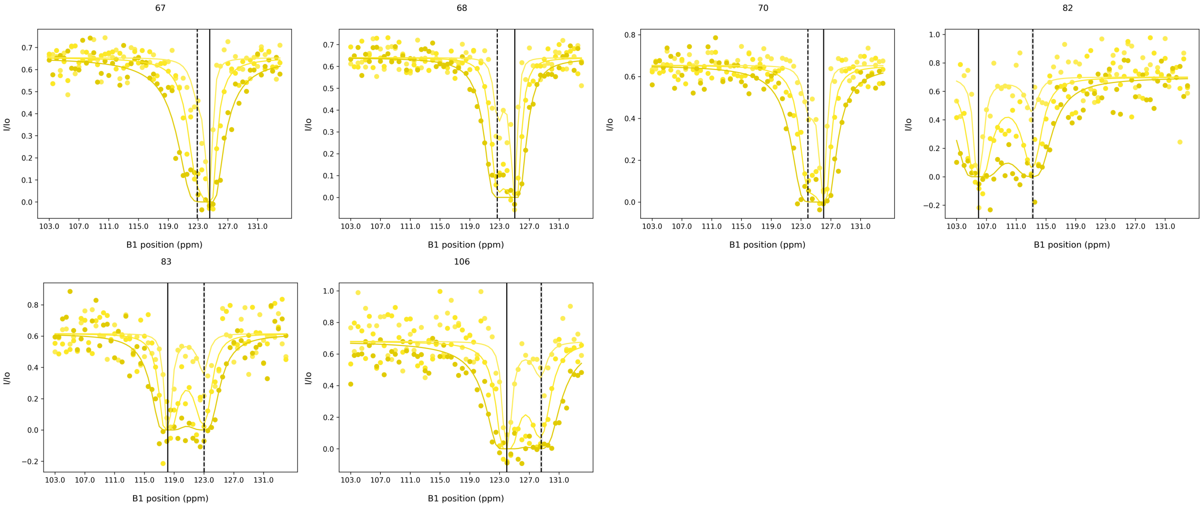

**Fig. 3.** CEST traces for 1 mM $CaCl_2$ sample. Data(circles) and fits(lines) for 50, 25, and 10 hz saturation field strengths. Solid vertical lines indicate the $^{15}N$ chemical shift for the major state, dotted vertical lines indicate the $^{15}N$ chemical shift for the minor state.

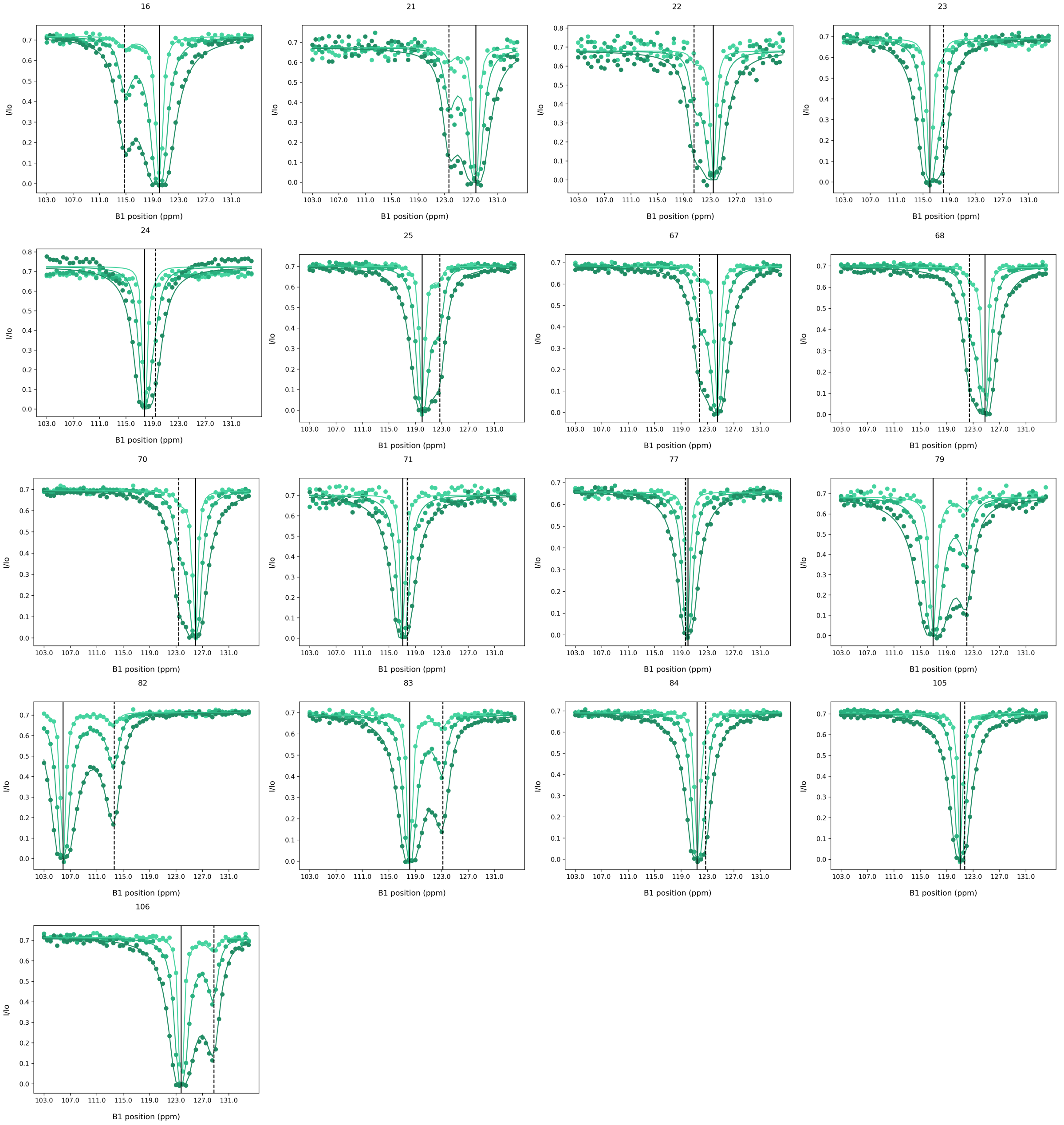

**Fig. 4.** CEST traces for 3 mM $CaCl_2$ sample. Data(circles) and fits(lines) for 50, 25, and 10 hz saturation field strengths. Solid vertical lines indicate the $^{15}N$ chemical shift for the major state, dotted vertical lines indicate the $^{15}N$ chemical shift for the minor state.

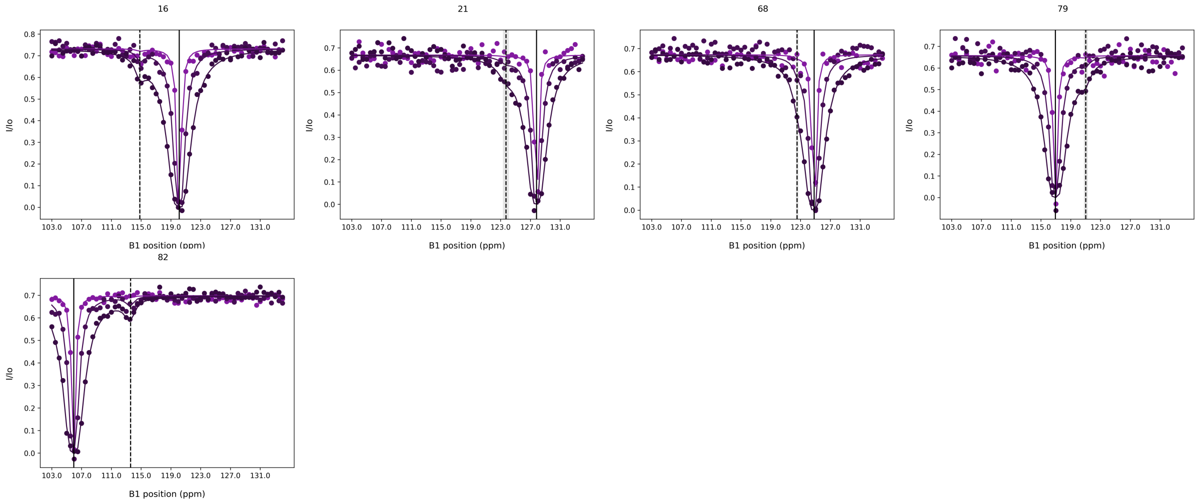

**Fig. 5.** CEST traces for 5 mM $CaCl_2$ sample. Data(circles) and fits(lines) for 50, 25, and 10 hz saturation field strengths. Solid vertical lines indicate the $^{15}N$ chemical shift for the major state, dotted vertical lines indicate the $^{15}N$ chemical shift for the minor state.

**Table 1.** FAST-Modelfree-derived parameters for residues at 5 mM $CaCl_2$

| 5 mM $Ca^{2+}$ | | | | | | | | | | |
|---|---|---|---|---|---|---|---|---|---|---|
| Residue | Model | $S^2$ | $S^2$ err | $S^2$f | $S^2$ferr | $\tau_e$ | $\tau_e$ err | $R_{ex}$ | $R_{ex}$ err | SSE |
| 1 | 1 | 0.806 | 0.026 | | | | | | | 14.52 |
| 3 | 1 | 0.976 | 0.017 | | | | | | | 0.527 |
| 5 | 1 | 0.995 | 0.009 | | | | | | | 0 |
| 6 | 1 | 0.968 | 0.016 | | | | | | | 3.559 |
| 7 | 1 | 0.957 | 0.03 | | | | | | | 1.573 |
| 8 | 1 | 0.976 | 0.023 | | | | | | | 12.81 |
| 9 | 3 | 0.931 | 0.035 | | | | | 2.642 | 0.727 | 0.376 |
| 10 | 3 | 0.934 | 0.019 | | | | | 2.867 | 0.352 | 0.049 |
| 11 | 1 | 0.965 | 0.022 | | | | | | | 1.857 |
| 12 | 1 | 1 | 0.01 | | | | | | | 0.45 |
| 13 | 3 | 0.914 | 0.059 | | | | | 5.253 | 1.039 | 0.088 |
| 14 | 1 | 0.979 | 0.02 | | | | | | | 0.39 |
| 16 | 3 | 0.819 | 0.057 | | | | | 9.409 | 1.141 | 2.968 |
| 18 | 1 | 0.974 | 0.03 | | | | | | | 3.75 |
| 19 | 3 | 0.963 | 0.063 | | | | | 10.939 | 3.141 | 0.353 |
| 20 | 3 | 0.957 | 0.031 | | | | | 7.257 | 0.886 | 1.033 |
| 21 | 1 | 0.964 | 0.03 | | | | | | | 5.205 |
| 22 | 1 | 0.979 | 0.022 | | | | | | | 3.93 |
| 23 | 1 | 1 | 0.025 | | | | | | | 2.561 |
| 24 | 1 | 0.934 | 0.024 | | | | | | | 7.42 |
| 25 | 1 | 0.963 | 0.027 | | | | | | | 3.701 |
| 27 | 1 | 0.917 | 0.033 | | | | | | | 1.627 |
| 28 | 1 | 0.998 | 0.019 | | | | | | | 0.048 |
| 29 | 1 | 0.998 | 0.021 | | | | | | | 0.86 |
| 30 | 1 | 0.987 | 0.023 | | | | | | | 5.478 |
| 31 | 1 | 0.929 | 0.042 | | | | | | | 1.203 |
| 32 | 1 | 0.936 | 0.027 | | | | | | | 10.51 |
| 33 | 1 | 0.968 | 0.018 | | | | | | | 0.41 |
| 34 | 1 | 0.949 | 0.02 | | | | | | | 3.324 |
| 35 | 1 | 0.881 | 0.038 | | | | | | | 2.25 |
| 36 | 1 | 0.987 | 0.028 | | | | | | | 1.477 |
| 37 | 1 | 0.962 | 0.016 | | | | | | | 1.559 |
| 38 | 1 | 1 | 0.013 | | | | | | | 0.838 |
| 39 | 1 | 0.979 | 0.014 | | | | | | | 10.86 |
| 40 | 1 | 0.912 | 0.047 | | | | | | | 0.994 |
| 41 | 1 | 0.95 | 0.046 | | | | | | | 3.295 |
| 42 | 1 | 0.922 | 0.023 | | | | | | | 3.881 |
| 43 | 1 | 0.956 | 0.029 | | | | | | | 3.707 |
| 45 | 3 | 0.761 | 0.052 | | | | | 4.096 | 1.151 | 1.104 |
| 46 | 1 | 0.954 | 0.024 | | | | | | | 0.737 |
| 47 | 1 | 0.953 | 0.024 | | | | | | | 1.071 |
| 48 | 1 | 0.998 | 0.026 | | | | | | | 0.419 |
| 49 | 1 | 0.889 | 0.025 | | | | | | | 1.209 |
| 50 | 1 | 0.904 | 0.021 | | | | | | | 4.171 |
| 51 | 1 | 0.948 | 0.033 | | | | | | | 1.747 |
| 52 | 1 | 0.847 | 0.026 | | | | | | | 1.189 |
| 53 | 1 | 0.943 | 0.012 | | | | | | | 8.879 |
| 54 | 1 | 0.923 | 0.03 | | | | | | | 0.818 |
| 55 | 1 | 1 | 0.017 | | | | | | | 8.062 |
| 56 | 1 | 1 | 0.033 | | | | | | | 0.392 |
| 57 | 1 | 0.879 | 0.026 | | | | | | | 8.75 |

Continued on next page

DRAFT

Continuation of Table 1

| Residue | Model | $S^2$ | $S^2$ err | $S^2$f | $S^2$ferr | $\tau_e$ | $\tau_e$ err | $R_{ex}$ | $R_{ex}$ err | SSE |
|---|---|---|---|---|---|---|---|---|---|---|
| 59 | 1 | 0.868 | 0.039 | | | | | | | 4.721 |
| 60 | 1 | 0.865 | 0.024 | | | | | | | 9.333 |
| 62 | 1 | 1 | 0.023 | | | | | | | 0.078 |
| 63 | 1 | 1 | 0.009 | | | | | | | 6.265 |
| 64 | 1 | 0.826 | 0.03 | | | | | | | 2.984 |
| 65 | 1 | 0.96 | 0.02 | | | | | | | 6.21 |
| 66 | 1 | 0.85 | 0.035 | | | | | | | 0.971 |
| 67 | 1 | 1 | 0.02 | | | | | | | 1.587 |
| 68 | 1 | 1 | 0.023 | | | | | | | 0.979 |
| 70 | 3 | 0.918 | 0.029 | | | | | 3.395 | 0.51 | 0.078 |
| 71 | 1 | 0.971 | 0.03 | | | | | | | 3.022 |
| 72 | 3 | 1 | 0.026 | | | | | 9.512 | 1.425 | 1.242 |
| 73 | 3 | 0.863 | 0.052 | | | | | 9.928 | 1.46 | 0.028 |
| 77 | 1 | 0.836 | 0.032 | | | | | | | 7.288 |
| 79 | 1 | 1 | 0.032 | | | | | | | 2.263 |
| 81 | 1 | 0.815 | 0.057 | | | | | | | 1.728 |
| 82 | 3 | 0.855 | 0.032 | | | | | 2.885 | 0.561 | 1.74 |
| 83 | 1 | 0.987 | 0.031 | | | | | | | 2.858 |
| 84 | 1 | 0.988 | 0.011 | | | | | | | 1.493 |
| 85 | 1 | 0.877 | 0.011 | | | | | | | 2.662 |
| 86 | 1 | 0.905 | 0.032 | | | | | | | 0.714 |
| 88 | 3 | 1 | 0.012 | | | | | 4.274 | 0.736 | 1.598 |
| 89 | 1 | 1 | 0.019 | | | | | | | 0.306 |
| 90 | 1 | 1 | 0.022 | | | | | | | 1.742 |
| 91 | 2 | 0.957 | 0.017 | | | | | | | 0.283 |
| 92 | 1 | 1 | 0.016 | | | | | | | 19.56 |
| 93 | 3 | 0.849 | 0.018 | | | | | 4.298 | 0.455 | 3.659 |
| 94 | 1 | 0.811 | 0.014 | | | | | | | 18.02 |
| 97 | 1 | 1 | 0.022 | | | | | | | 3.85 |
| 98 | 3 | 0.934 | 0.056 | | | | | 18.216 | 3.778 | 2.02 |
| 99 | 3 | 1 | 0.012 | | | | | 3.162 | 0.467 | 0.418 |
| 100 | 1 | 0.906 | 0.027 | | | | | | | 4.843 |
| 101 | 1 | 0.874 | 0.026 | | | | | | | 1.198 |
| 102 | 1 | 0.856 | 0.02 | | | | | | | 0.314 |
| 103 | 1 | 0.931 | 0.021 | | | | | | | 1.856 |
| 105 | 1 | 0.924 | 0.031 | | | | | | | 0.462 |
| 107 | 1 | 0.961 | 0.035 | | | | | | | 2.64 |
| 108 | 3 | 0.709 | 0.065 | | | | | 4.617 | 1.394 | 0.228 |
| 110 | 1 | 0.971 | 0.014 | | | | | | | 0.366 |
| 111 | 1 | 0.965 | 0.02 | | | | | | | 0.681 |
| 113 | 1 | 0.967 | 0.024 | | | | | | | 19.36 |
| 114 | 1 | 1 | 0.019 | | | | | | | 3.585 |
| 115 | 1 | 1 | 0.012 | | | | | | | 3.741 |
| 116 | 1 | 1 | 0.023 | | | | | | | 4.785 |
| 117 | 1 | 1 | 0.015 | | | | | | | 1.747 |
| 118 | 1 | 1 | 0.024 | | | | | | | 2.971 |
| 119 | 1 | 0.967 | 0.037 | | | | | | | 1.867 |
| 120 | 1 | 0.945 | 0.04 | | | | | | | 2.791 |
| 121 | 1 | 0.972 | 0.036 | | | | | | | 0.536 |
| 122 | 1 | 0.989 | 0.023 | | | | | | | 6.327 |

End of Table

DRAFT

**Table 2.** FAST-Modelfree-derived parameters for residues at 3 mM $CaCl_2$

| 3 mM $Ca^{2+}$ | | | | | | | | | | |
|---|---|---|---|---|---|---|---|---|---|---|
| Residue | Model | $S^2$ | $S^2$ err | $S^2f$ | $S^2ferr$ | $\tau_e$ | $\tau_e$ err | $R_{ex}$ | $R_{ex}$ err | SSE |
| 1 | 1 | 0.825 | 0.052 | | | | | | | 4.265 |
| 3 | 1 | 0.888 | 0.024 | | | | | | | 0.992 |
| 4 | 1 | 0.828 | 0.021 | | | | | | | 4.44 |
| 5 | 1 | 0.921 | 0.019 | | | | | | | 2.144 |
| 6 | 1 | 0.896 | 0.036 | | | | | | | 2.514 |
| 7 | 1 | 0.982 | 0.011 | | | | | | | 5.556 |
| 8 | 1 | 0.83 | 0.016 | | | | | | | 5.233 |
| 9 | 1 | 0.958 | 0.027 | | | | | | | 5.49 |
| 10 | 3 | 0.834 | 0.046 | | | | | 2.851 | 0.99 | 0.206 |
| 11 | 1 | 0.933 | 0.028 | | | | | | | 3.771 |
| 12 | 1 | 0.992 | 0.017 | | | | | | | 3.065 |
| 13 | 3 | 0.839 | 0.056 | | | | | 13.11 | 1.524 | 0.699 |
| 14 | 1 | 1 | 0.034 | | | | | | | 3.804 |
| 18 | 1 | 0.908 | 0.061 | | | | | | | 0.226 |
| 19 | 3 | 0.859 | 0.102 | | | | | 35.082 | 7.337 | 0.216 |
| 20 | 3 | 0.945 | 0.067 | | | | | 7.854 | 2.167 | 0.129 |
| 21 | 3 | 0.878 | 0.087 | | | | | 5.134 | 1.781 | 0.124 |
| 22 | 1 | 0.943 | 0.046 | | | | | | | 5.57 |
| 23 | 1 | 1 | 0.026 | | | | | | | 5.733 |
| 24 | 1 | 0.97 | 0.018 | | | | | | | 1.683 |
| 25 | 1 | 0.913 | 0.027 | | | | | | | 0.656 |
| 26 | 1 | 1 | 0.014 | | | | | | | 2.234 |
| 27 | 1 | 0.874 | 0.013 | | | | | | | 8.33 |
| 28 | 1 | 0.909 | 0.029 | | | | | | | 0.643 |
| 29 | 1 | 0.78 | 0.062 | | | | | | | 1.479 |
| 30 | 1 | 0.784 | 0.032 | | | | | | | 4.448 |
| 31 | 1 | 0.884 | 0.021 | | | | | | | 2.892 |
| 32 | 1 | 0.885 | 0.04 | | | | | | | 2.282 |
| 33 | 1 | 0.859 | 0.032 | | | | | | | 4.551 |
| 34 | 1 | 0.844 | 0.034 | | | | | | | 0.514 |
| 35 | 1 | 0.907 | 0.024 | | | | | | | 0.115 |
| 36 | 3 | 0.873 | 0.038 | | | | | 2.447 | 0.632 | 0.453 |
| 37 | 1 | 0.945 | 0.013 | | | | | | | 1.765 |
| 38 | 1 | 1 | 0.022 | | | | | | | 6.27 |
| 39 | 1 | 0.969 | 0.032 | | | | | | | 5.227 |
| 40 | 1 | 1 | 0.03 | | | | | | | 3.413 |
| 41 | 1 | 0.916 | 0.047 | | | | | | | 0.969 |
| 42 | 1 | 0.882 | 0.023 | | | | | | | 1.961 |
| 45 | 3 | 0.747 | 0.041 | | | | | 4.232 | 1.224 | 0.135 |
| 46 | 1 | 0.893 | 0.02 | | | | | | | 5.35 |
| 47 | 1 | 0.901 | 0.014 | | | | | | | 1.239 |
| 50 | 1 | 0.824 | 0.021 | | | | | | | 4.386 |
| 51 | 1 | 0.836 | 0.052 | | | | | | | 0.552 |
| 52 | 1 | 0.86 | 0.022 | | | | | | | 1.139 |
| 53 | 1 | 0.737 | 0.049 | | | | | | | 1.722 |
| 54 | 1 | 0.829 | 0.031 | | | | | | | 1.056 |
| 55 | 1 | 0.989 | 0.023 | | | | | | | 1.94 |
| 56 | 1 | 1 | 0.05 | | | | | | | 2.806 |
| 57 | 1 | 0.81 | 0.021 | | | | | | | 15.43 |
| 59 | 1 | 0.77 | 0.035 | | | | | | | 2.233 |
| 62 | 1 | 0.946 | 0.069 | | | | | | | 0.413 |

Continued on next page

DRAFT

Continuation of Table 2

| Residue | Model | $S^2$ | $S^2$ err | $S^2$f | $S^2$ferr | $\tau_e$ | $\tau_e$ err | $R_{ex}$ | $R_{ex}$ err | SSE |
|---|---|---|---|---|---|---|---|---|---|---|
| 63 | 1 | 0.896 | 0.068 | | | | | | | 1.262 |
| 64 | 1 | 0.79 | 0.06 | | | | | | | 0.737 |
| 65 | 1 | 0.81 | 0.02 | | | | | | | 12.46 |
| 66 | 1 | 0.906 | 0.019 | | | | | | | 0.824 |
| 67 | 3 | 0.932 | 0.031 | | | | | 2.724 | 0.694 | 0.076 |
| 68 | 3 | 0.831 | 0.045 | | | | | 3.375 | 0.78 | 0.288 |
| 69 | 1 | 0.925 | 0.023 | | | | | | | 0.457 |
| 70 | 1 | 0.959 | 0.03 | | | | | | | 4.173 |
| 71 | 1 | 0.987 | 0.031 | | | | | | | 2.854 |
| 72 | 1 | 1 | 0.09 | | | | | | | 4.047 |
| 77 | 1 | 0.837 | 0.062 | | | | | | | 4.732 |
| 82 | 3 | 0.85 | 0.054 | | | | | 5.696 | 1.015 | 2.099 |
| 83 | 3 | 0.866 | 0.04 | | | | | 5.954 | 0.995 | 0.078 |
| 84 | 3 | 0.918 | 0.042 | | | | | 2.18 | 0.669 | 0.257 |
| 85 | 1 | 0.823 | 0.017 | | | | | | | 0.348 |
| 86 | 1 | 0.864 | 0.01 | | | | | | | 1.887 |
| 88 | 1 | 1 | 0.016 | | | | | | | 2.192 |
| 89 | 1 | 1 | 0.018 | | | | | | | 0.145 |
| 91 | 1 | 0.921 | 0.016 | | | | | | | 13.91 |
| 92 | 1 | 0.985 | 0.018 | | | | | | | 7.877 |
| 93 | 4 | 0.759 | 0.028 | | | 19.538 | 6.593 | 3.557 | 0.515 | 0 |
| 94 | 5 | 0.624 | 0.04 | 0.748 | 0.034 | 1075.8 | 372.08 | | | 0 |
| 97 | 1 | 1 | 0.02 | | | | | | | 0.04 |
| 98 | 3 | 0.987 | 0.076 | | | | | 32.222 | 4.224 | 0.128 |
| 99 | 1 | 1 | 0.015 | | | | | | | 3.306 |
| 100 | 1 | 0.819 | 0.024 | | | | | | | 3.633 |
| 101 | 1 | 0.886 | 0.022 | | | | | | | 0.53 |
| 102 | 1 | 0.782 | 0.053 | | | | | | | 0.767 |
| 103 | 1 | 0.833 | 0.023 | | | | | | | 5.713 |
| 105 | 1 | 0.905 | 0.016 | | | | | | | 0.736 |
| 106 | 3 | 0.853 | 0.033 | | | | | 5.926 | 0.96 | 1.556 |
| 107 | 1 | 0.896 | 0.022 | | | | | | | 0.585 |
| 108 | 1 | 0.843 | 0.06 | | | | | | | 0.964 |
| 110 | 1 | 0.883 | 0.023 | | | | | | | 9.192 |
| 111 | 1 | 0.892 | 0.014 | | | | | | | 3.407 |
| 113 | 3 | 0.737 | 0.039 | | | | | 4.363 | 0.737 | 0.05 |
| 114 | 1 | 0.973 | 0.032 | | | | | | | 4.981 |
| 115 | 1 | 0.969 | 0.02 | | | | | | | 1.478 |
| 116 | 1 | 0.956 | 0.018 | | | | | | | 6.304 |
| 117 | 1 | 0.944 | 0.017 | | | | | | | 1.917 |
| 118 | 3 | 0.886 | 0.024 | | | | | 1.347 | 0.494 | 0.941 |
| 119 | 1 | 0.955 | 0.014 | | | | | | | 0.401 |
| 120 | 1 | 0.913 | 0.023 | | | | | | | 1.538 |
| 121 | 1 | 0.96 | 0.013 | | | | | | | 2.369 |
| 122 | 1 | 0.959 | 0.017 | | | | | | | 4.746 |
| 118 | 1 | 1 | 0.024 | | | | | | | 2.971 |
| 119 | 1 | 0.967 | 0.037 | | | | | | | 1.867 |
| 120 | 1 | 0.945 | 0.04 | | | | | | | 2.791 |
| 121 | 1 | 0.972 | 0.036 | | | | | | | 0.536 |
| 122 | 1 | 0.989 | 0.023 | | | | | | | 6.327 |

End of Table

DRAFT

**Table 3.** FAST-Modelfree-derived parameters for residues at 1 mM $CaCl_2$

| 1 mM $Ca^{2+}$ | | | | | | | | | | |
|---|---|---|---|---|---|---|---|---|---|---|
| Residue | Model | $S^2$ | $S^2$ err | $S^2f$ | $S^2$ferr | $\tau_e$ | $\tau_e$ err | $R_{ex}$ | $R_{ex}$ err | SSE |
| 1 | 5 | 0.435 | 0.026 | 0.66 | 0.01 | 2004.3 | 1167.4 | | | 0 |
| 2 | 1 | 0.938 | 0.001 | | | | | | | 4.533 |
| 3 | 3 | 0.895 | 0.011 | | | | | 0.546 | 0.154 | 0.072 |
| 4 | 1 | 0.893 | 0.001 | | | | | | | 0.133 |
| 5 | 1 | 0.948 | 0.002 | | | | | | | 5.207 |
| 6 | 3 | 0.896 | 0.02 | | | | | 4.053 | 0.298 | 0.215 |
| 7 | 1 | 0.904 | 0.002 | | | | | | | 10.75 |
| 8 | 3 | 0.82 | 0.013 | | | | | 4.376 | 0.21 | 0.955 |
| 9 | 3 | 0.889 | 0.058 | | | | | 4.17 | 0.836 | 0.007 |
| 10 | 3 | 0.847 | 0.029 | | | | | 5.236 | 0.416 | 1.23 |
| 11 | 1 | 0.914 | 0.004 | | | | | | | 0.207 |
| 12 | 1 | 0.823 | 0.011 | | | | | | | 6.583 |
| 22 | 3 | 0.884 | 0.089 | | | | | 4.528 | 1.3 | 0.014 |
| 25 | 3 | 0.869 | 0.025 | | | | | 3.532 | 0.373 | 0.615 |
| 26 | 3 | 0.901 | 0.034 | | | | | 2.572 | 0.495 | 0.127 |
| 27 | 2 | 0.842 | 0.001 | | | 40.67 | 8.547 | | | 2.997 |
| 29 | 1 | 0.819 | 0.004 | | | | | | | 15.68 |
| 32 | 1 | 0.894 | 0.002 | | | | | | | 14.58 |
| 33 | 2 | 0.86 | 0.002 | | | 36.888 | 10.745 | | | 2.682 |
| 34 | 3 | 0.86 | 0.01 | | | | | 1.726 | 0.137 | 0.084 |
| 35 | 1 | 0.862 | 0.008 | | | | | | | 5.696 |
| 36 | 3 | 0.855 | 0.019 | | | | | 6.388 | 0.279 | 2.485 |
| 37 | 1 | 1 | 0.002 | | | | | | | 1.139 |
| 38 | 3 | 1 | 0.12 | | | | | 14.826 | 2.089 | 0.504 |
| 39 | 3 | 0.913 | 0.021 | | | | | 3.267 | 0.307 | 0.435 |
| 41 | 3 | 0.79 | 0.073 | | | | | 3.1 | 1.061 | 0.143 |
| 43 | 3 | 0.858 | 0.027 | | | | | 1.523 | 0.398 | 1.778 |
| 45 | 3 | 0.715 | 0.061 | | | | | 7.458 | 0.901 | 0.246 |
| 46 | 3 | 0.906 | 0.011 | | | | | 0.727 | 0.158 | 1.328 |
| 47 | 1 | 0.94 | 0.004 | | | | | | | 2.045 |
| 48 | 3 | 0.962 | 0.024 | | | | | 0.913 | 0.34 | 0.111 |
| 49 | 1 | 0.855 | 0.002 | | | | | | | 6.501 |
| 50 | 1 | 0.849 | 0.002 | | | | | | | 11.95 |
| 51 | 1 | 0.866 | 0.003 | | | | | | | 0.724 |
| 52 | 1 | 0.895 | 0.001 | | | | | | | 0.246 |
| 53 | 3 | 0.82 | 0.01 | | | | | 1.089 | 0.145 | 0.35 |
| 54 | 1 | 0.885 | 0.002 | | | | | | | 2.031 |
| 55 | 2 | 0.965 | 0.003 | | | 1866.9 | 378.38 | | | 1.147 |
| 56 | 3 | 0.98 | 0.033 | | | | | 1.999 | 0.478 | 0.085 |
| 59 | 1 | 0.885 | 0.003 | | | | | | | 4.86 |
| 62 | 1 | 0.903 | 0.004 | | | | | | | 1.787 |
| 63 | 1 | 0.964 | 0.001 | | | | | | | 2.879 |
| 64 | 1 | 0.843 | 0.003 | | | | | | | 15.01 |
| 65 | 1 | 0.85 | 0.002 | | | | | | | 12.44 |
| 66 | 3 | 0.884 | 0.024 | | | | | 2.188 | 0.363 | 0.016 |
| 67 | 3 | 0.935 | 0.04 | | | | | 4.591 | 0.585 | 1 |
| 82 | 3 | 0.933 | 0.071 | | | | | 19.875 | 1.041 | 0.059 |
| 83 | 3 | 0.874 | 0.052 | | | | | 10.91 | 0.776 | 1.397 |
| 85 | 1 | 0.86 | 0.002 | | | | | | | 3.61 |
| 86 | 1 | 0.917 | 0.002 | | | | | | | 2.545 |
| 88 | 4 | 0.923 | 0.017 | | | 96.11 | 33.974 | 4.045 | 0.235 | 0 |

Continued on next page

DRAFT

Continuation of Table 3

| Residue | Model | $S^2$ | $S^2$ err | $S^2$f | $S^2$ferr | $\tau_e$ | $\tau_e$ err | $R_{ex}$ | $R_{ex}$ err | SSE |
|---|---|---|---|---|---|---|---|---|---|---|
| 89 | 1 | 0.938 | 0.003 | | | | | | | 0.703 |
| 90 | 3 | 0.975 | 0.011 | | | | | 0.445 | 0.158 | 0.08 |
| 91 | 1 | 0.896 | 0.002 | | | | | | | 10.26 |
| 92 | 3 | 0.874 | 0.014 | | | | | 2.338 | 0.212 | 0.021 |
| 93 | 3 | 0.811 | 0.008 | | | | | 3.137 | 0.121 | 2.455 |
| 94 | 5 | 0.698 | 0.003 | 0.789 | 0.007 | 1092.9 | 201.65 | | | 0 |
| 97 | 1 | 0.984 | 0.001 | | | | | | | 12.79 |
| 98 | 3 | 0.92 | 0.034 | | | | | 33.159 | 0.506 | 0.019 |
| 99 | 1 | 1 | 0.001 | | | | | | | 13.21 |
| 100 | 1 | 0.82 | 0.001 | | | | | | | 3.247 |
| 101 | 1 | 0.906 | 0.002 | | | | | | | 0.875 |
| 102 | 1 | 0.871 | 0.003 | | | | | | | 12.46 |
| 103 | 1 | 0.791 | 0.001 | | | | | | | 8.769 |
| 105 | 3 | 0.878 | 0.028 | | | | | 2.456 | 0.411 | 1.643 |
| 106 | 3 | 0.803 | 0.05 | | | | | 19.15 | 0.736 | 0.486 |
| 107 | 1 | 0.951 | 0.004 | | | | | | | 6.617 |
| 108 | 1 | 0.878 | 0.004 | | | | | | | 3.513 |
| 111 | 3 | 0.896 | 0.008 | | | | | 0.674 | 0.124 | 0.003 |
| 114 | 1 | 0.968 | 0.001 | | | | | | | 18.56 |
| 115 | 1 | 0.938 | 0.004 | | | | | | | 8.241 |
| 116 | 3 | 0.872 | 0.011 | | | | | 2.2 | 0.163 | 0.078 |
| 117 | 3 | 0.899 | 0.01 | | | | | 3.18 | 0.145 | 0.088 |
| 118 | 3 | 0.832 | 0.013 | | | | | 4.1 | 0.189 | 0.114 |
| 119 | 1 | 1 | 0.001 | | | | | | | 13.27 |
| 120 | 1 | 1 | 0.001 | | | | | | | 1.98 |
| 121 | 3 | 0.924 | 0.017 | | | | | 1.432 | 0.245 | 0.932 |
| 122 | 1 | 0.977 | 0.002 | | | | | | | 2.749 |
| 102 | 1 | 0.782 | 0.053 | | | | | | | 0.767 |
| 103 | 1 | 0.833 | 0.023 | | | | | | | 5.713 |
| 105 | 1 | 0.905 | 0.016 | | | | | | | 0.736 |
| 106 | 3 | 0.853 | 0.033 | | | | | 5.926 | 0.96 | 1.556 |
| 107 | 1 | 0.896 | 0.022 | | | | | | | 0.585 |
| 108 | 1 | 0.843 | 0.06 | | | | | | | 0.964 |
| 110 | 1 | 0.883 | 0.023 | | | | | | | 9.192 |
| 111 | 1 | 0.892 | 0.014 | | | | | | | 3.407 |
| 113 | 3 | 0.737 | 0.039 | | | | | 4.363 | 0.737 | 0.05 |
| 114 | 1 | 0.973 | 0.032 | | | | | | | 4.981 |
| 115 | 1 | 0.969 | 0.02 | | | | | | | 1.478 |
| 116 | 1 | 0.956 | 0.018 | | | | | | | 6.304 |
| 117 | 1 | 0.944 | 0.017 | | | | | | | 1.917 |
| 118 | 3 | 0.886 | 0.024 | | | | | 1.347 | 0.494 | 0.941 |
| 119 | 1 | 0.955 | 0.014 | | | | | | | 0.401 |
| 120 | 1 | 0.913 | 0.023 | | | | | | | 1.538 |
| 121 | 1 | 0.96 | 0.013 | | | | | | | 2.369 |
| 122 | 1 | 0.959 | 0.017 | | | | | | | 4.746 |

End of Table

DRAFT

**Table 4.** FAST-Modelfree-derived parameters for residues at 5 mM $CaCl_2$ and Saturating (7.6:1) IP3

| 5 mM $Ca^{2+}$ and Saturating (7.6:1) IP3 | | | | | | | | | | |
|---|---|---|---|---|---|---|---|---|---|---|
| Residue | Model | $S^2$ | $S^2$ err | $S^2f$ | $S^2ferr$ | $\tau_e$ | $\tau_e$ err | $R_{ex}$ | $R_{ex}$ err | SSE |
| 1 | 1 | 0.867 | 0.017 | | | | | | | 1.259 |
| 2 | 1 | 0.923 | 0.018 | | | | | | | 7.612 |
| 3 | 1 | 0.896 | 0.018 | | | | | | | 2.442 |
| 4 | 1 | 0.863 | 0.014 | | | | | | | 4.144 |
| 5 | 1 | 0.872 | 0.013 | | | | | | | 9.321 |
| 6 | 1 | 0.886 | 0.014 | | | | | | | 3.981 |
| 7 | 1 | 0.909 | 0.018 | | | | | | | 10.06 |
| 8 | 3 | 0.834 | 0.011 | | | | | 0.837 | 0.298 | 0.383 |
| 9 | 1 | 0.908 | 0.014 | | | | | | | 2.149 |
| 10 | 1 | 0.885 | 0.014 | | | | | | | 17.51 |
| 11 | 1 | 0.868 | 0.016 | | | | | | | 1.477 |
| 12 | 1 | 0.907 | 0.011 | | | | | | | 0.943 |
| 13 | 3 | 0.952 | 0.026 | | | | | 2.892 | 0.595 | 0.598 |
| 14 | 1 | 0.81 | 0.013 | | | | | | | 0.557 |
| 16 | 3 | 0.758 | 0.019 | | | | | 6.499 | 0.376 | 0.86 |
| 18 | 1 | 0.924 | 0.022 | | | | | | | 4.528 |
| 20 | 3 | 0.819 | 0.023 | | | | | 3.412 | 0.532 | 0.083 |
| 21 | 1 | 0.85 | 0.017 | | | | | | | 0.572 |
| 22 | 1 | 0.851 | 0.027 | | | | | | | 0.941 |
| 24 | 3 | 0.878 | 0.022 | | | | | 1.752 | 0.394 | 1.41 |
| 25 | 1 | 0.896 | 0.009 | | | | | | | 6.326 |
| 26 | 1 | 0.938 | 0.018 | | | | | | | 9.71 |
| 27 | 1 | 0.826 | 0.009 | | | | | | | 16.77 |
| 28 | 1 | 0.871 | 0.011 | | | | | | | 4.451 |
| 29 | 1 | 0.833 | 0.022 | | | | | | | 9.223 |
| 30 | 1 | 0.86 | 0.015 | | | | | | | 2.276 |
| 31 | 4 | 0.754 | 0.026 | | | 54.477 | 16.279 | 3.24 | 0.477 | 0 |
| 32 | 2 | 0.813 | 0.014 | | | 95.817 | 25.081 | | | 3.061 |
| 33 | 1 | 0.889 | 0.018 | | | | | | | 3.213 |
| 34 | 1 | 0.84 | 0.01 | | | | | | | 0.142 |
| 35 | 1 | 0.882 | 0.013 | | | | | | | 10.95 |
| 36 | 3 | 0.859 | 0.019 | | | | | 1.8 | 0.39 | 0.827 |
| 37 | 1 | 0.872 | 0.009 | | | | | | | 3.21 |
| 38 | 1 | 0.895 | 0.021 | | | | | | | 6.176 |
| 39 | 1 | 0.855 | 0.011 | | | | | | | 0.751 |
| 41 | 1 | 0.859 | 0.019 | | | | | | | 0.101 |
| 42 | 1 | 0.871 | 0.015 | | | | | | | 5.412 |
| 43 | 1 | 0.853 | 0.014 | | | | | | | 17.49 |
| 45 | 3 | 0.705 | 0.018 | | | | | 2.533 | 0.472 | 1.9 |
| 46 | 1 | 0.841 | 0.016 | | | | | | | 0.31 |
| 47 | 1 | 0.84 | 0.011 | | | | | | | 10.27 |
| 48 | 1 | 0.906 | 0.012 | | | | | | | 3.218 |
| 49 | 2 | 0.815 | 0.017 | | | 40.763 | 13.993 | | | 0.636 |
| 50 | 1 | 0.845 | 0.012 | | | | | | | 0.775 |
| 51 | 1 | 0.869 | 0.018 | | | | | | | 4.356 |
| 52 | 1 | 0.841 | 0.016 | | | | | | | 0.597 |
| 53 | 1 | 0.826 | 0.008 | | | | | | | 10.31 |
| 54 | 1 | 0.899 | 0.018 | | | | | | | 2.748 |
| 55 | 1 | 0.984 | 0.021 | | | | | | | 0.115 |
| 56 | 1 | 0.962 | 0.033 | | | | | | | 4.073 |
| 59 | 1 | 0.781 | 0.027 | | | | | | | 1.61 |

Continued on next page

DRAFT

Continuation of Table 4

| Residue | Model | $S^2$ | $S^2$ err | $S^2f$ | $S^2ferr$ | $\tau_e$ | $\tau_e$ err | $R_{ex}$ | $R_{ex}$ err | SSE |
|---|---|---|---|---|---|---|---|---|---|---|
| 60 | 5 | 0.724 | 0.03 | 0.919 | 0.028 | 500 | 65.06 | | | 0 |
| 62 | 1 | 0.918 | 0.028 | | | | | | | 6.445 |
| 63 | 1 | 0.886 | 0.014 | | | | | | | 3.76 |
| 66 | 3 | 0.845 | 0.018 | | | | | 1.466 | 0.458 | 0.229 |
| 67 | 1 | 0.949 | 0.012 | | | | | | | 0.787 |
| 68 | 1 | 0.884 | 0.017 | | | | | | | 5.134 |
| 69 | 1 | 0.901 | 0.013 | | | | | | | 0.315 |
| 70 | 1 | 0.931 | 0.015 | | | | | | | 4.713 |
| 71 | 1 | 0.891 | 0.014 | | | | | | | 3.634 |
| 72 | 1 | 0.967 | 0.017 | | | | | | | 0.861 |
| 73 | 3 | 0.806 | 0.025 | | | | | 2.532 | 0.53 | 0.326 |
| 77 | 1 | 0.857 | 0.012 | | | | | | | 1.551 |
| 79 | 1 | 1 | 0.017 | | | | | | | 2.633 |
| 81 | 2 | 0.854 | 0.015 | | | 40.164 | 15.146 | | | 0.005 |
| 82 | 1 | 0.848 | 0.011 | | | | | | | 2.556 |
| 83 | 1 | 0.954 | 0.016 | | | | | | | 8.284 |
| 84 | 1 | 0.856 | 0.012 | | | | | | | 2.966 |
| 85 | 1 | 0.797 | 0.006 | | | | | | | 3.594 |
| 86 | 2 | 0.823 | 0.016 | | | 59.158 | 16.71 | | | 0.888 |
| 88 | 3 | 0.929 | 0.023 | | | | | 2.433 | 0.614 | 2.028 |
| 89 | 1 | 0.949 | 0.016 | | | | | | | 2.292 |
| 90 | 2 | 0.889 | 0.029 | | | 989.46 | 397.99 | 0.902 | | |
| 92 | 1 | 0.93 | 0.018 | | | | | | | 2.289 |
| 93 | 4 | 0.738 | 0.016 | | | 23.504 | 7.673 | 3.466 | 0.281 | 0 |
| 99 | 3 | 0.901 | 0.02 | | | | | 1.977 | 0.444 | 0.932 |
| 100 | 2 | 0.786 | 0.009 | | | 31.475 | 7.619 | | | 2.187 |
| 101 | 1 | 0.844 | 0.014 | | | | | | | 0.655 |
| 102 | 1 | 0.785 | 0.011 | | | | | | | 4.402 |
| 103 | 1 | 0.823 | 0.011 | | | | | | | 4.297 |
| 105 | 1 | 0.877 | 0.009 | | | | | | | 0.443 |
| 106 | 1 | 0.879 | 0.02 | | | | | | | 2.049 |
| 107 | 1 | 0.861 | 0.012 | | | | | | | 1.764 |
| 108 | 1 | 0.883 | 0.022 | | | | | | | 1.378 |
| 109 | 1 | 0.849 | 0.009 | | | | | | | 4.562 |
| 110 | 1 | 0.886 | 0.009 | | | | | | | 4.798 |
| 111 | 1 | 0.906 | 0.011 | | | | | | | 5.13 |
| 113 | 3 | 0.805 | 0.015 | | | | | 1.564 | 0.545 | 0 |
| 114 | 1 | 0.909 | 0.012 | | | | | | | 3.885 |
| 116 | 3 | 0.875 | 0.011 | | | | | 1.125 | 0.288 | 0.193 |
| 117 | 1 | 0.886 | 0.011 | | | | | | | 1.21 |
| 118 | 3 | 0.803 | 0.015 | | | | | 1.899 | 0.506 | 0.111 |
| 119 | 1 | 0.923 | 0.012 | | | | | | | 0.168 |
| 120 | 1 | 0.861 | 0.012 | | | | | | | 5.366 |
| 121 | 1 | 0.898 | 0.015 | | | | | | | 2.247 |
| 122 | 3 | 0.858 | 0.019 | | | | | 1.085 | 0.38 | 0.042 |

End of Table

DRAFT

**Table 5.** CEST-derived chemical shifts for exchanging residues at 1 mM $CaCl_2$

| 1 mM $Ca^{2+}$ | | |
|---|---|---|
| Residue | Chemical Shift State A | Chemical Shift State B |
| 67 | 124.56 ± 0.05 | 122.90 ± 0.12 |
| 68 | 125.05 ± 0.05 | 122.69 ± 0.09 |
| 70 | 126.03 ± 0.04 | 123.92 ± 0.10 |
| 82 | 105.94 ± 0.10 | 113.23 ± 0.18 |
| 83 | 118.13 ± 0.10 | 123.03 ± 0.17 |
| 84 | 122.21 ± 0.04 | 121.09 ± 0.06 |
| 106 | 123.98 ± 0.07 | 128.60 ± 0.18 |

DRAFT

**Table 6.** CEST-derived chemical shifts for exchanging residues at 3 mM $CaCl_2$

| 3 mM $Ca^{2+}$ | | |
|---|---|---|
| Residue | Chemical Shift State A | Chemical Shift State B |
| 16 | 120.06 ± 0.01 | 114.77 ± 0.03 |
| 21 | 127.76 ± 0.02 | 123.68 ± 0.07 |
| 22 | 123.48 ± 0.03 | 120.56 ± 0.08 |
| 23 | 116.04 ± 0.01 | 118.13 ± 0.03 |
| 24 | 117.85 ± 0.01 | 119.46 ± 0.04 |
| 25 | 120.03 ± 0.01 | 122.72 ± 0.01 |
| 67 | 124.52 ± 0.01 | 121.82 ± 0.02 |
| 68 | 124.80 ± 0.01 | 122.43 ± 0.02 |
| 70 | 125.95 ± 0.01 | 123.40 ± 0.02 |
| 71 | 117.09 ± 0.02 | 117.80 ± 0.15 |
| 77 | 120.06 ± 0.02 | 119.69 ± 0.16 |
| 79 | 116.95 ± 0.02 | 122.05 ± 0.04 |
| 82 | 105.90 ± 0.01 | 113.63 ± 0.01 |
| 83 | 118.16 ± 0.01 | 123.17 ± 0.02 |
| 84 | 121.14 ± 0.01 | 122.74 ± 0.02 |
| 105 | 121.05 ± 0.01 | 121.74 ± 0.04 |
| 106 | 123.78 ± 0.01 | 128.75 ± 0.02 |

DRAFT

**Table 7.** CEST-derived chemical shifts for exchanging residues at 5 mM $CaCl_2$

| 5 mM $Ca^{2+}$ | | |
|---|---|---|
| Residue | Chemical Shift State A | Chemical Shift State B |
| 16 | 120.13 ±0.01 | 114.83 ± 0.01 |
| 21 | 127.83 ± 0.01 | 123.72 ± 0.04 |
| 68 | 124.83 ± 0.01 | 122.54 ± 0.02 |
| 79 | 116.94 ± 0.03 | 121.03 ± 0.03 |
| 82 | 105.95 ± 0.01 | 113.59 ± 0.01 |

DRAFT
